# Differential Regulation of Single Microtubules and Bundles by a Three-Protein Module

**DOI:** 10.1101/2020.03.05.979864

**Authors:** Nandini Mani, Shuo Jiang, Alex E. Neary, Sithara S. Wijeratne, Radhika Subramanian

## Abstract

A remarkable feature of the microtubule cytoskeleton is co-existence of sub-populations having different dynamic properties. A prominent example is the anaphase spindle, where stable antiparallel bundles exist alongside dynamic microtubules and provide spatial cues for cytokinesis. How are dynamics of spatially proximal arrays differentially regulated? We reconstitute a minimal system of three midzone proteins: microtubule-crosslinker PRC1, and its interactors CLASP1 and Kif4A, proteins that promote and suppress microtubule elongation, respectively. We find their collective activity promotes elongation of single microtubules, while simultaneously stalling polymerization of crosslinked bundles. This differentiation arises from (i) Strong rescue activity of CLASP1, which overcomes weaker effects of Kif4A on single microtubules, (ii) Lower microtubule and PRC1-binding affinity of CLASP1, which permit dominance of Kif4A at overlaps. In addition to canonical mechanisms where antagonistic regulators set microtubule lengths, our findings illuminate design principles by which collective regulator activity creates microenvironments of arrays with distinct dynamic properties.

## INTRODUCTION

Microtubules form the backbone of micron-sized structures such as mitotic spindles, plant cortical arrays and neuronal axons. A remarkable feature of these structures is the co-existence of microtubule sub-populations with distinct dynamic properties, each tuned to specific functions^1,2^. For instance, axons contain long stable microtubules that form tracks for intra-cellular transport and exist alongside short dynamic microtubules that serve as microtubule seeds and sources of tubulin^3^. In the plant cortex, microtubule bundles are organized into stable arrays that specify the axis of cell elongation^4^. However, the simultaneous presence of dynamic microtubules in the cortex is important in reorienting arrays in response to environmental signals. How the dynamics of distinct microtubule populations present in close proximity are differentially regulated is a fundamental question.

Differential regulation of dynamics is prominent among the various sub-structures of the mitotic spindle such as interpolar, midzone, astral and kinetochore microtubules^5-8^. During the metaphase to anaphase transition, spindle microtubules are reorganized to form a crosslinked, antiparallel array at the cell center known as the spindle midzone^9^. This structure keeps separating chromosomes apart and provides positional cues for cell cleavage. Antiparallel microtubule arrays at the cell center are stable with low tubulin turnover frequencies^10,11^. In addition, the cell center also contains single dynamic microtubules arising from nucleation and polymerization of new microtubules in the inter-polar region of the cell^12-14^. How newly nucleated microtubules continue to grow while the crosslinked bundles present alongside them are largely stabilized is unclear.

The dynamics of microtubule populations at the spindle midzone are regulated by several Microtubule Associated Proteins (MAPs). In particular, the localization of three classes of MAPs to the cell center is conserved: (i) cross-linkers of antiparallel microtubules such as Protein Regulator of Cytokinesis-1 (PRC1)^15-19^ (ii) suppressors of microtubule growth such as kinesins Kif4A or Kip3p^20-22^, and (iii) growth promoters, which are most often homologues of mammalian CLASP1^23-25^. In this study, we focus on the collective activity of the mammalian PRC1-Kif4A-CLASP1 module in regulating the dynamics of different microtubule sub-populations.

PRC1 is a conserved non-motor MAP that specifically cross-links microtubules in an antiparallel orientation^26,27^. PRC1 directly binds and recruits other proteins to the cell center, including CLASP1 and Kif4A^28-31^. During anaphase, CLASP1 contributes to initiating the assembly of microtubules in the central spindle, and is important for their growth and stability^23-25,32,33^, while Kif4A regulates the length of microtubules in the spindle midzone^20,28^. *In vitro*, CLASP1 and Kif4A can each autonomously bind microtubules and regulate dynamics. CLASP1 is a rescue factor that suppresses microtubule catastrophes and promotes rescues, to increase the lengths of dynamic microtubules. While Kif4A also inhibits catastrophes, it reduces microtubule growth rate to limit polymer length. Microtubules are less dynamic in the presence of Kif4A due to the reduction of tubulin turnover at their plus-ends^34-40^. The individual activities of CLASP1 and Kif4A thus have opposite outcomes on the lengths and dynamicity of microtubules, with CLASP1 promoting elongation and Kif4A suppressing growth. The regulation of microtubule dynamics by the collective activity of CLASP1 and Kif4A has not been studied either in the context of single microtubules or antiparallel bundles.

A largely unexplored question in regulating the architecture of complex microtubule networks with different subpopulations such as the spindle midzone, is how can the dynamics of distinct spatially proximal microtubule arrays be differentially regulated? An intuitive mechanism is to spatially segregate MAPs among subsets of microtubules^41^. However, segregation is difficult at the cell center where dynamic microtubule tips and stable overlaps occur within 0.6-0.7 μm of each other^14^, and microtubule-MAP interactions are typically characterized by micromolar binding affinities. Moreover, this strategy does not permit individual arrays to switch between stable and dynamic states. An alternate mechanism that would allow for such switching is to modulate MAP activity on specific arrays through post-translational modifications. However, when MAPs are present at high cytosolic concentrations and turnover rapidly between microtubule-bound and soluble fractions, it is challenging to regulate only a subset of MAP molecules through post-translational modifications. Overall, it is unclear if the dynamics of proximal microtubule arrays can be differentially regulated simply by the collective activity of a set of MAPs in the absence of external factors such as regulatory proteins. Further, it is not known how they can do so in a manner that allows individual arrays to switch between dynamic states, which would be essential in forming large adaptive microtubule networks such as the spindle.

Here we report that a minimal protein module comprising of three spindle midzone proteins, PRC1, Kif4A and CLASP1, promotes the elongation of single microtubules, while simultaneously suppressing the growth of crosslinked microtubules. Central to this is an “inverse microtubule affinity-activity relationship” for each regulator, with high rescue activity-low microtubule affinity of CLASP1 molecules and low growth suppression activity-high microtubule affinity of Kif4A motors, along with different PRC1-binding affinities. Our findings illuminate the design principles underlying differential regulation of proximal microtubule arrays by the collective activity of MAPs that have opposite effects on microtubule length and dynamicity.

## RESULTS

### CLASP1 dominates over Kif4A on single microtubules

We first determined the collective effect of Kif4A and CLASP1 on the length-regulation of single microtubules in the absence of PRC1. Recombinant CLASP1-GFP and Kif4A were purified from Sf9 cells (Extended data Fig. 1a). SEC-MALS (Size Exclusion Chromatography coupled with Multi-Angle Light Scattering) analysis of three constructs of CLASP1 that included different subsets of domains tagged with GFP (1-654, 654-1471 and 805-1471) showed that they are all monomeric in solution (Extended data Fig. 1b). Dynamic light scattering studies further showed that CLASP1(805-1471)-GFP had low polydispersity and was elongated in shape (Extended data Fig. 1c). Full-length CLASP1 is thus likely to be a monomer in solution^42^.

The dynamics of single microtubules in the presence of full-length CLASP1 and Kif4A was visualized using multi-wavelength TIRF microscopy (Extended data Fig. 2a). In controls with tubulin alone, microtubules exhibited dynamic instability, where rescue events were rare and most catastrophe events result in depolymerization of the microtubule back to the seed (hereafter referred to as “complete catastrophe”) (Fig. 1a). In experiments with 200 nM CLASP1-GFP alone, a dramatic reduction of complete catastrophes was observed. With 1000nM Kif4A alone, we saw a striking reduction in microtubule growth from seeds and an increase in duration of pauses where no polymerization was recorded, consistent with known activities of these proteins^35,36,38^ (Fig. 1a).

**Figure 1:**
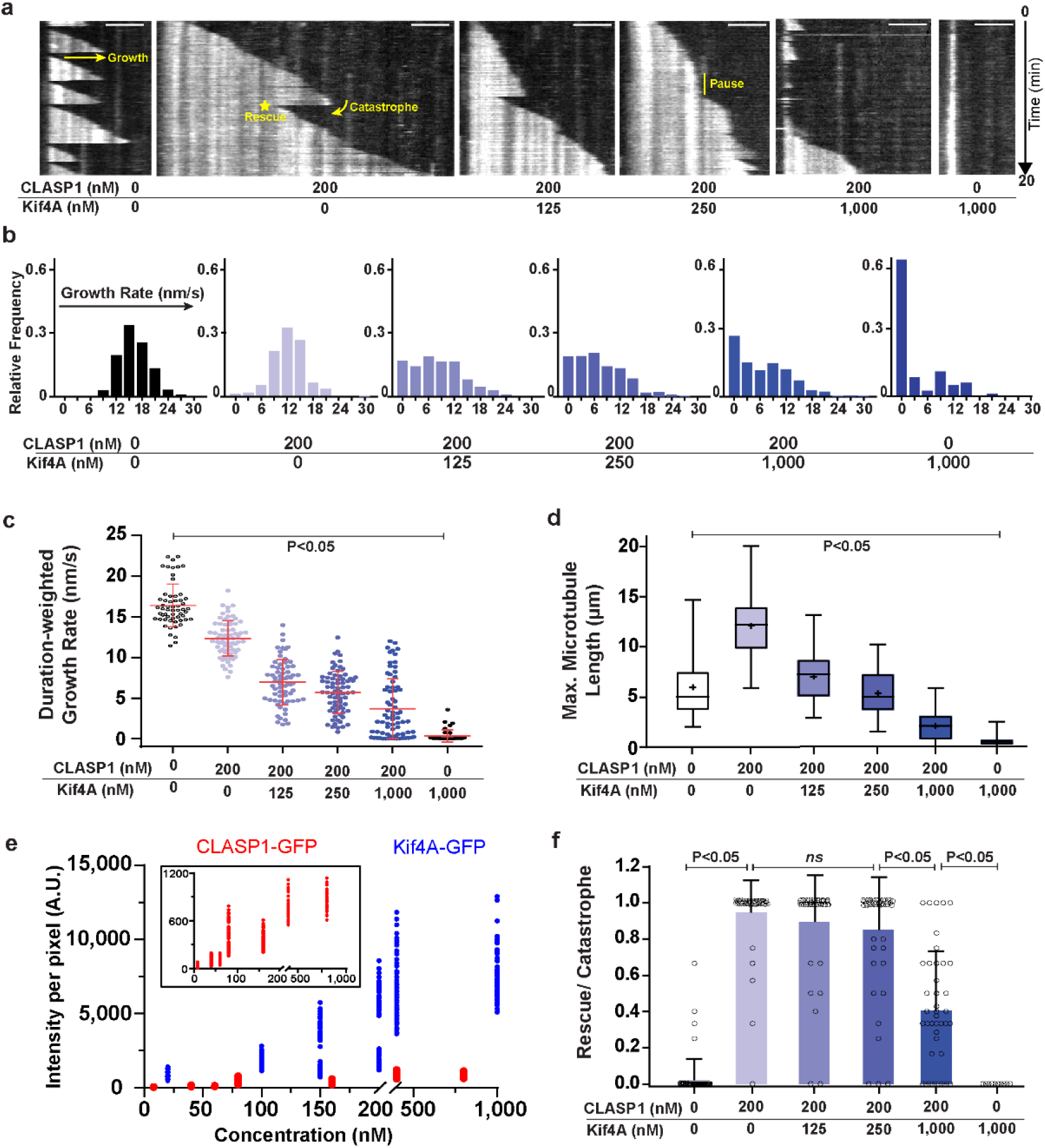
The rescue activity of CLASP1 overrides growth suppression by Kif4A on single microtubules. Also see Extended data Figs. 1, 2 and 3 n = total number of kymographs analyzed from a total of 3 independent experiments for each condition, except where indicated. All concentrations correspond to dimeric Kif4A and monomeric CLASP1-GFP, except where indicated. P-values were calculated from an ordinary one-way ANOVA test with Dunnett correction for multiple comparisons. **a.** Representative kymographs of X-rhodamine-microtubules in the presence of Kif4A and CLASP1-GFP. GMPCPP-seeds are not shown. **→** indicates growth event and direction of growth. Curved arrow indicates a catastrophe event. * indicates a rescue event on the microtubule. | indicates a period of stalled growth (‘pause’) on the microtubule. Scale bar represents 2 μm. Number of kymographs examined: tubulin control (65), 200 nM CLASP1 (76), 200 nM CLASP1 + 125 nM Kif4A (69), 200 nM CLASP1 + 250 nM Kif4A (73), 200 nM CLASP1 + 1000 nM Kif4A (73) and 1000 nM Kif4A (38). **b.** Histogram of growth rates for each condition seen in (**a**). Number of events analyzed: tubulin control (306), 200 nM CLASP1 (218), 200 nM CLASP1 + 125 nM Kif4A (308), 200 nM CLASP1 + 250 nM Kif4A (381), 200 nM CLASP1 + 1000 nM Kif4A (331) and 1000 nM Kif4A (78). **c.** Scatter plot of duration-weighted microtubule growth rate. Mean (center line) and standard deviation as indicated by red bars, for assay conditions: tubulin control (16.4 ± 2.6 nm/s, n = 58), 200 nM CLASP1 (12.4 ± 2.2 nm/s, n = 66), 200 nM CLASP1 + 125 nM Kif4A (6.9 ± 2.8 nm/s, n = 67), 200 nM CLASP1 + 250 nM Kif4A (5.7 ± 2.6 nm/s, n = 73), 200 nM CLASP1 + 1000 nM Kif4A (3.7 ± 3.7 nm/s, n = 73) and 1000 nM Kif4A (0.3 ± 0.8 nm/s, n = 38). P < 0.0001 for each column when compared to tubulin alone. **d.** Box and whisker plot of maximum microtubule length. Horizontal lines within box indicate the 25^th^, median (center line) and 75^th^ percentile. Plus-sign indicates mean. Error bars indicate minimum and maximum range. Mean and standard deviation for assay conditions with tubulin alone (6.0 ± 2.9 μm, n = 60), 200 nM CLASP1 (12.1 ± 3.2 μm, n = 68), 200 nM CLASP1 + 125 nM Kif4A (7.0 ± 2.3 μm, n = 67), 200 nM CLASP1 + 250 nM Kif4A (5.4 ± 2.1 μm, n = 72), 200 nM CLASP1 + 1000 nM Kif4A (2.2 ± 1.4 μm, n = 74) and 1000 nM Kif4A (0.6 ± 0.5 μm, n = 40). P < 0.0001 for 1000 nM Kif4A when compared to the tubulin control. **e.** Scatter plot of GFP intensity per pixel on taxol-stabilized microtubules in the presence of Kif4A-GFP (•) or CLASP1-GFP (•). All concentrations refer to *monomeric* proteins. Inset shows magnified region containing intensities for CLASP1-GFP(•). Mean and standard deviation of intensities for assay conditions with CLASP1-GFP: 8 nM (26.9 ±18, n=70), 40 nM (95.3 ± 41.8, n=70), 60 nM (109 ± 30.0, n=70), 80 nM (400.1 ± 166.9, n=70), 160 nM (397.6 ± 90.3, n=70), 400 nM (754.5 ± 154.3, n=70), 800 nM (854.3 ± 122.1, n=35). Assay conditions with Kif4A-GFP: 20 nM (894.0 ± 253.9, n=70), 100 nM (1856 ± 380.3, n=70), 150 nM (2649 ± 1527, n=70), 200 nM (3982 ± 2346, n=70), 400 nM (6814 ± 2085, n=70), 1000 nM (7542 ± 1640, n=70). n refers to number of microtubules analyzed from 2 independent experiments for each condition. **f.** Scatter plot and Bar graph showing mean of the ratio of rescue to total number of catastrophe events in 20 minutes. Error bars indicate standard deviation. Assay conditions: Tubulin control (0.0 ± 0.1, n = 60), 200 nM CLASP1 (1.0 ± 0.2, n = 57), 200 nM CLASP1 + 125 nM Kif4A (0.9 ± 0.3, n = 43), 200 nM CLASP1 + 250 nM Kif4A (0.9 ± 0.3, n = 49), 200 nM CLASP1 + 1000 nM Kif4A (0.4 ± 0.3, n = 42) and 1000 nM Kif4A (0.0 ± 0.0, n = 10). P < 0.0001 for tubulin control to 200 nM CLASP1, for 200 nM CLASP1 + 250 nM Kif4A to 200 nM CLASP1 + 1000 nM Kif4A and to 1000 nM Kif4A. P is not significant for 200 nM CLASP1 to (i) 200 nM CLASP1 + 125 nM Kif4A and to (ii) 200 nM CLASP1 + 250 nM Kif4A.

We next held the CLASP1-GFP concentration constant (200 nM) and titrated in increasing amounts of Kif4A (125 - 1000 nM). Addition of increasing amounts of Kif4A resulted in lowering of polymerization rates and an increase in the frequency of the pause-state where polymerization is stalled (Fig. 1a-1b). In addition to pause frequency, pause duration also increased (Extended data Fig. 2b). We computed a “duration-weighted growth rate” for each kymograph to quantify growth rate while accounting for differences in duration of each phase (Methods, Fig. 1c). This parameter decreased systematically with increasing ratios of Kif4A:CLASP1, consistent with lower growth rate and increased duration of pausing at higher Kif4A concentrations.

We next examined how the combined activities of these proteins regulates microtubule length. Compared to tubulin-control, experiments with 200 nM CLASP1-GFP showed that the maximum length a microtubule grew from the seeds in the 20 min observation time, and the average microtubule length before any catastrophe event increased 2-fold (Fig. 1d, Extended data Fig. 2c). Addition of increasing amounts of Kif4A along with 200 nM CLASP1-GFP systematically decreased the maximum microtubule length (Fig. 1d). We noticed that the tubulin control and experiments with 200 nM CLASP1-GFP + 250 nM Kif4A resulted in similar maximum microtubule lengths, even though the duration-weighted growth rate dropped by a factor of ~3 (Fig. 1c). The maximum microtubule length became lower than the tubulin-control only with the addition of 1000 nM Kif4A (Fig. 1d). This suggests that the activity of CLASP1 in promoting the elongation of microtubules dominates over the growth-suppressing activity of Kif4A and nearly 5-fold excess of Kif4A over CLASP1 is needed before Kif4A can effectively suppress microtubule elongation.

### How does CLASP1 counteract growth suppression by Kif4A?

The ability of CLASP1 to override Kif4A activity on length regulation of single microtubules could arise due to either (i) higher microtubule-binding of CLASP1 compared to Kif4A, or (ii) suppression of catastrophes by CLASP1, or (iii) promotion of rescues by CLASP1.

We compared microtubule-binding of Kif4A and CLASP1, through fluorescence intensity analysis of respective GFP-tagged proteins on stabilized microtubules. The experiments were performed at low ATP (150 nM), where Kif4A binds uniformly along the microtubule lattice. The fluorescence intensity/pixel of Kif4A-GFP on microtubules was 4-9 fold higher than CLASP1-GFP (Fig. 1e). Similar experiments with higher ATP concentrations excluded the possibility that low concentrations of ATP promoted stronger Kif4A-GFP binding (Extended data Fig. 2d). These results indicate that Kif4A has a stronger affinity for microtubule-binding than CLASP1. We next compared the affinity of each protein for the ends of dynamic microtubules by performing single molecule experiments with CLASP1-GFP and Kif4A-GFP. Single molecules of CLASP1 bound and diffused all along the microtubule lattice, whereas Kif4A molecules moved processively towards and accumulated at the plus-end (Extended data fig. 3a-b). Ratio of tip to lattice fluorescence intensities showed that the plus-end occupancy of Kif4A was higher than CLASP1 under conditions where the total microtubule-bound intensities of both proteins were similar (Extended data Figs. 3c-d). Therefore, the elongation of microtubules in mixtures of CLASP1 and Kif4A does not arise from a higher microtubule-occupancy of CLASP1 relative to Kif4A.

Next, we considered if the increased microtubule elongation could arise from suppression of catastrophe by Kif4A and CLASP1. However, under all assay conditions, the catastrophe frequency was uniformly ~ 2-2.5 fold less than control (Extended data Fig. 2e). Finally, we examined rescue frequency and found it increased 20-fold with 200 nM CLASP1-GFP compared to tubulin-control. No rescue events were observed with 1000 nM Kif4A alone. However, in all Kif4A:CLASP1 ratios tested, the rescue frequency remained higher than tubulin-control, with nearly 100% of all catastrophe events being rescued by 200 nM CLASP1-GFP (Fig. 1f, Extended data Fig. 2f). Remarkably, even with a 5-fold excess of Kif4A over CLASP1, nearly 40% of catastrophe events are rescued, despite a systematic decrease in the levels of microtubule-bound CLASP1-GFP with increasing Kif4A (Extended data Fig. 2g).

Together the data suggest that CLASP1’s activity as a rescue-promoting factor dominates on single microtubules over a wide concentration range, counteracting growth suppression by Kif4A to promote polymer elongation. Remarkably, this activity of CLASP1 dominates despite its weaker microtubule-binding affinity relative to Kif4A.

### Differential regulation of single and crosslinked microtubules

In addition to binding to microtubules, both Kif4A and CLASP1 are known to bind to PRC1^28,29^, raising the question if dynamics of PRC1-crosslinked microtubules are similar to those of single microtubules in the presence of Kif4A and CLASP1.

To answer this, we reconstituted the collective activity of Kif4A, CLASP1 and PRC1 in assays where we can simultaneously visualize the dynamics of both crosslinked and single microtubules (Fig. 2a, hereafter referred to as “dynamic bundles assay”). In these experiments we can observe the elongation of preformed antiparallel microtubule overlaps and the formation of new overlaps from cross-linking of growing microtubules. In the presence of 0.5 nM PRC1 and 5 nM Kif4A, microtubule dynamics were completely suppressed, and microtubules exhibited relative sliding^43,44^ (Extended data Fig. 4a). With 0.5 nM PRC1 and 200 nM CLASP1-GFP, kymographs and montages reveal a near-continuous extension of microtubules from plus-ends of both single microtubules and preformed overlaps for the duration of the experiment indicating that CLASP1 is active on both sets of microtubules under this condition (Figs. 2b-2c, Extended data Fig. 4b, Supplementary video 1). The addition of 10 nM Kif4A along with 0.5 nM PRC1 and 200 nM CLASP1-GFP resulted in continuous elongation of single microtubules, as they did in the absence of Kif4A. However, crosslinked microtubules in the same field of view showed a dramatic suppression of microtubule polymerization (Figs. 2d-2e, Extended data Fig. 4c, Supplementary video 2). Thus, a minimal system consisting of three proteins can differentially regulate the dynamics of single and crosslinked microtubules.

**Figure 2:**
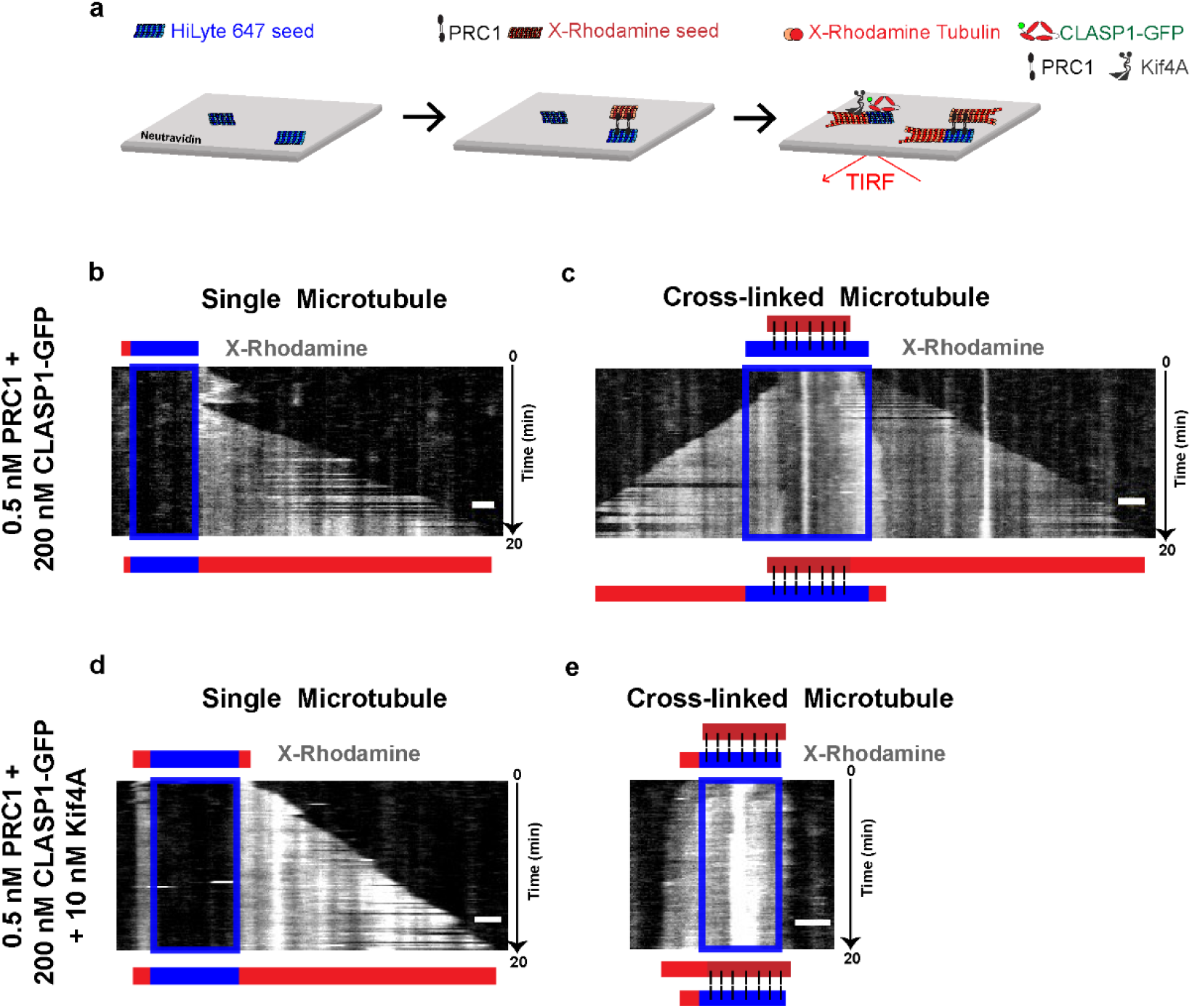
The collective activity of Kif4A, CLASP1 and PRC1 differentially regulates the dynamics of single and cross-linked microtubules. Also see Extended data Fig. 4, Supplementary Video 1 and Supplementary Video 2. **a.** Schematic of the dynamic microtubule assay used to examine the collective activity of PRC1, Kif4A and CLASP1-GFP on cross-linked and single microtubules. **b-e.** Representative kymograph of X-Rhodamine channel of single **(b,d)** and cross-linked microtubule **(c,e)**. The position of the seed is shown as a blue box. The schematics above and below the kymograph indicate the positions of the seed (blue) and microtubules (red) at the start and end of the experiment, respectively. Scale bar represents 2 μm. Assay conditions: **(b,c)** 0.5 nM PRC1 + 200 nM CLASP1-GFP (Kymographs of 47 single and 29 cross-linked microtubules were examined), **(d,e)** 0.5 nM PRC1 + 200 nM CLASP1-GFP + 10 nM Kif4A (Kymographs of 42 single and 49 cross-linked microtubules were examined).

### Quantitative analysis of dynamics of crosslinked microtubules

How does the collective activity of PRC1, Kif4A and CLASP1 regulate the dynamics of crosslinked overlaps? We systematically increased the Kif4A concentration from 10 – 125 nM while keeping CLASP1-GFP concentration constant at 200 nM and found that there was no microtubule elongation in over 70% of all preformed bundles analyzed (Extended data Fig. 5a). Instead, polymerization stalled when two anti-parallel microtubules formed a bundle and relative sliding was observed (Fig. 3a, Extended data Figs. 5b-5c). In contrast, at Kif4A concentrations below 10 nM, microtubule elongation was observed at all overlaps (Fig. 3b-3c, Extended data Fig. 5a).

**Figure 3:**
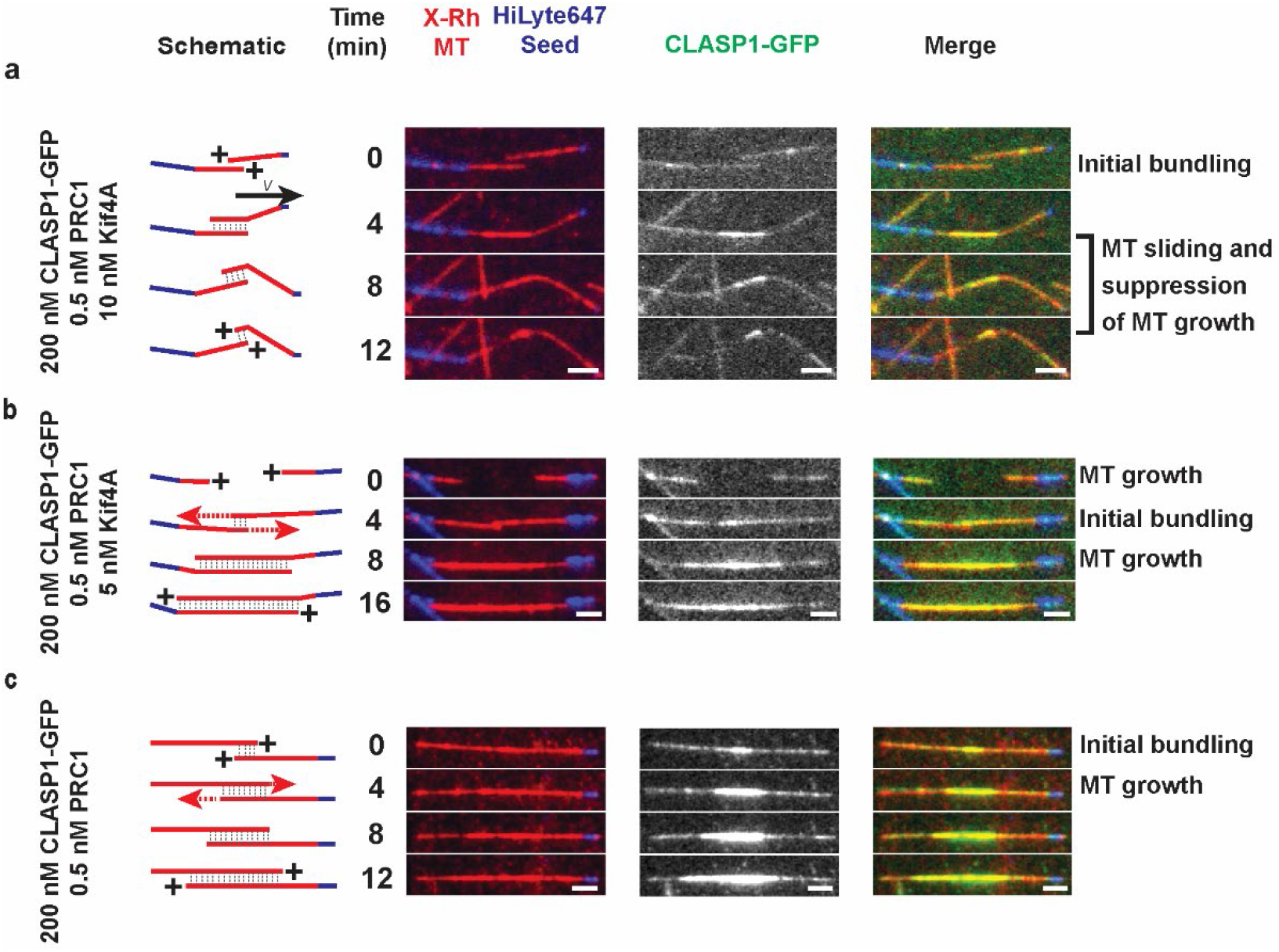
Dynamics of cross-linked microtubules in presence of Kif4A, CLASP1 and PRC1. Also see Extended data Fig. 5 Schematics and representative montages of microtubule bundles (red) grown from microtubule seeds (blue) in 3 independent experiments. Schematics indicate the plus end of the microtubules within the bundle. Velocity arrow indicates direction of microtubule sliding. Dotted red arrows indicate microtubule growth. Dotted gray lines indicate regions of overlap. X-Rh MT: X-Rhodamine microtubules. Scale bar represents 2 μm. Assay conditions and number of events (n) examined are **(a)** 200 nM CLASP1-GFP + 0.5 nM PRC1 + 10 nM Kif4A (n=36), **(b)** 200 nM CLASP1-GFP + 0.5 nM PRC1 + 5 nM Kif4A (n=37), **(c)** 200 nM CLASP1-GFP + 0.5 nM PRC1 (n = 33)

Consistent with this, duration-weighted growth rate for microtubule overlaps was close to 0 at Kif4A concentrations greater than 10 nM, and significant polymerization was apparent when Kif4A concentration is reduced to 5 nM (Fig. 4a). Similarly, plots of maximum microtubule length and average length of overlaps revealed no extension until the Kif4A concentration was reduced to 5 nM in the presence of 200 nM CLASP1-GFP and 0.5 nM PRC1 (Fig. 4b, Extended data Fig. 6a). Together, these data show that Kif4A has a dominant effect in regulating the dynamics of crosslinked microtubules over a wide concentration range.

**Figure 4:**
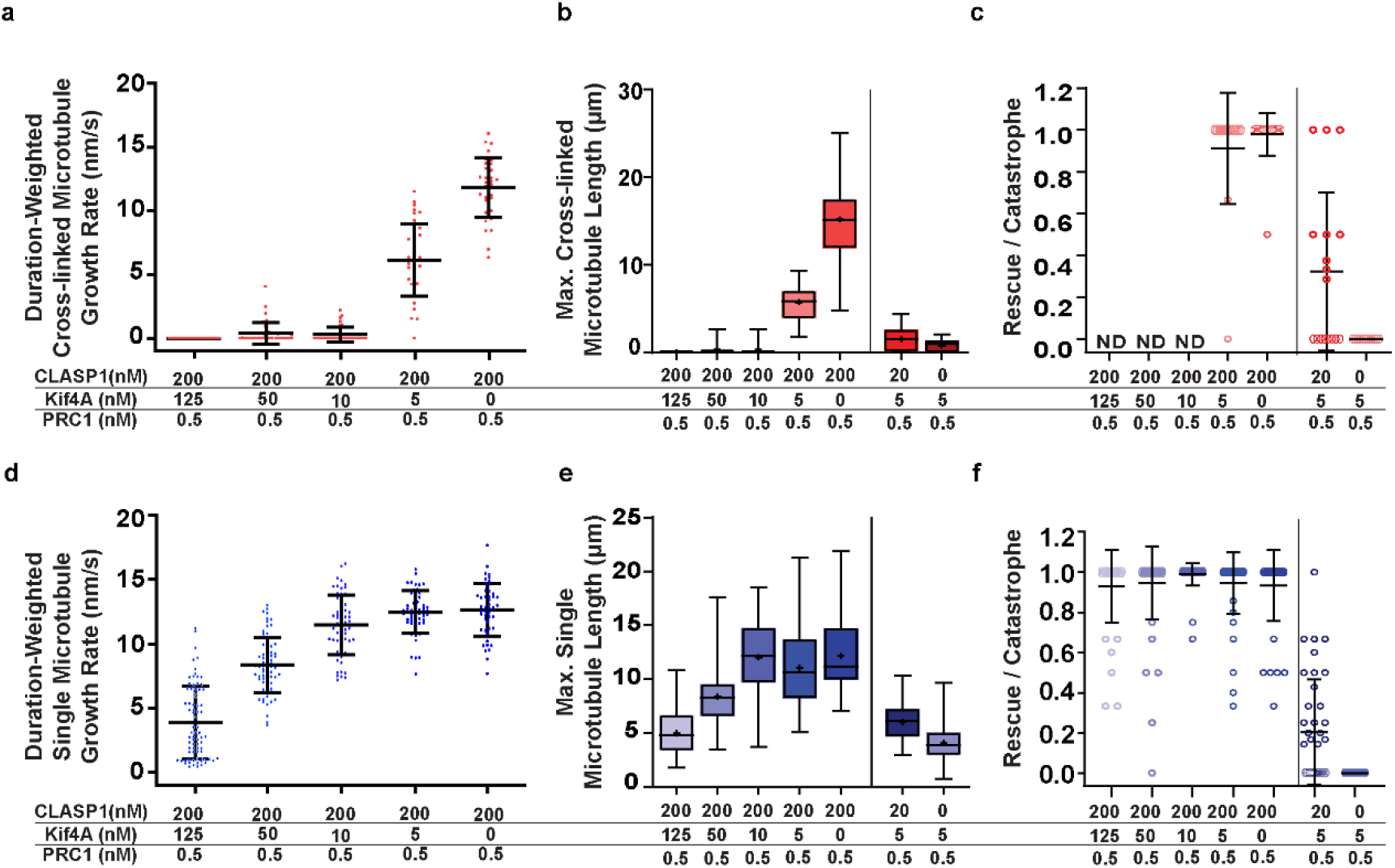
Kif4A activity dominates to suppress dynamics of cross-linked microtubules while CLASP1 activity promotes the elongation of single microtubules under identical reaction conditions. Also see Extended data Fig. 6 n = total number of kymographs of overlaps **(a-c)** and single microtubules **(d-f)** analyzed from 3 independent experiments in each condition **a.** Scatter plot of duration-weighted growth rate of cross-linked microtubules. Black bars indicate mean (center line) and standard deviation. Assay conditions: 200 nM CLASP1 + 0.5 nM PRC1 + 125 nM Kif4A 0.0 ± 0.0 nm/s [n = 30]; 200 nM CLASP1 + 0.5 nM PRC1 + 50 nM Kif4A 0.4 ± 0.8 nm/s [n = 39]; 200 nM CLASP1 + 0.5 nM PRC1 + 10 nM Kif4A 0.3 ± 0.6 nm/s [n = 35]; 200 nM CLASP1 + 0.5 nM PRC1 + 5 nM Kif4A 6.1 ± 2.8 nm/s [n = 36] and 200 nM + 0.5 nM PRC1 11.8 ± 2.3 nm/s [n = 33] **b.** Box and whisker plot of maximum length of cross-linked overlap. Plus-sign indicates mean. Horizontal lines within box indicate the 25^th^, median (center line) and 75^th^ percentile. Error bars indicate minimum and maximum range. Mean and standard deviation for assay conditions: 200 nM CLASP1 + 0.5 nM PRC1 + 125 nM Kif4A (0.0 ± 0.0 μm, n = 30), 200 nM CLASP1 + 0.5 nM PRC1 + 50 nM Kif4A (0.3 ± 0.6 μm, n = 33), 200 nM CLASP1 + 0.5 nM PRC1 + 10 nM Kif4A (0.3 ± 0.7 μm, n = 36), 200 nM CLASP1 + 0.5 nM PRC1 + 5 nM Kif4A (5.7 ± 2.1 μm, n = 37), 200 nM + 0.5 nM PRC1 (15.2 ± 4.4 μm, n = 33), 20 nM CLASP1 + 0.5 nM PRC1 + 5 nM Kif4A (1.5 ± 1.3 μm, n = 24) and 0.5 nM PRC1 + 5 nM Kif4A (0.8 ± 0.7 μm, n = 20). **c.** Scatter plot of the ratio of rescues to total number of catastrophe events in microtubule bundles in 20 minutes. Black bars indicate mean (center line) and standard deviation. ND: Not determined because no dynamics were observed. Assay conditions: 200 nM CLASP1 + 0.5 nM PRC1 + 125 nM Kif4A (*Not Determined*, n = 30), 200 nM CLASP1 + 0.5 nM PRC1 + 50 nM Kif4A (*Not Determined*, n = 39), 200 nM CLASP1 + 0.5 nM PRC1 + 10 nM Kif4A (*Not Determined*, n = 36), 200 nM CLASP1 + 0.5 nM PRC1 + 5 nM Kif4A (0.9 ± 0.3, n = 15), 200 nM + 0.5 nM PRC1 (1.0 ± 0.1, n = 24), 20 nM CLASP1 + 0.5 nM PRC1 + 5 nM Kif4A (0.3 ± 0.4, n = 17) and 0.5 nM PRC1 + 5 nM Kif4A (0.0 ± 0.0, n = 12). **d.** Scatter plot of duration-weighted growth rates of single microtubules. Black bars indicate mean (center line) and standard deviation. Assay conditions: 200 nM CLASP1 + 0.5 nM PRC1 + 125 nM Kif4A (3.9 ± 2.9 nm/s, n = 7); 200 nM CLASP1 + 0.5 nM PRC1 + 50 nM Kif4A (8.4 ± 2.2 nm/s, n = 69); 200 nM CLASP1 + 0.5 nM PRC1 + 10 nM Kif4A (11.3 ± 2.5 nm/s, n = 60); 200 nM CLASP1 + 0.5 nM PRC1 + 5 nM Kif4A (12.5 ± 1.7 nm/s, n = 52); and 200 nM + 0.5 nM PRC1 (12.7 ± 2.1 nm/s, n = 47). **e.** Box and whisker plot of maximum length of single microtubules. Plus-sign indicates mean. Horizontal lines within box indicate the 25^th^, median (center line) and 75^th^ percentile. Error bars indicate minimum and maximum range. Mean and standard deviation for assay conditions: 200 nM CLASP1 + 0.5 nM PRC1 + 125 nM Kif4A (5.0 ± 2.1 μm, n = 76); 200 nM CLASP1 + 0.5 nM PRC1 + 50 nM Kif4A (8.4 ± 2.8 μm, n = 69); 200 nM CLASP1 + 0.5 nM PRC1 + 10 nM Kif4A (12.1 ± 3.3 μm, n = 60); 200 nM CLASP1 + 0.5 nM PRC1 + 5 nM Kif4A (11.1 ± 3.8 μm, n = 53); and 200 nM + 0.5 nM PRC1 (12.2 ± 3.4 μm, n = 47). **f.** Bar graph of the ratio of rescues to total number of catastrophe events in single microtubules in 20 minutes. Black bars indicate mean (center line) and standard deviation. Assay conditions: 200 nM CLASP1 + 0.5 nM PRC1 + 125 nM Kif4A (0.93 ± 0.18, n = 41); 200 nM CLASP1 + 0.5 nM PRC1 + 50 nM Kif4A (0.95 ± 0.18, n = 63); 200 nM CLASP1 + 0.5 nM PRC1 + 10 nM Kif4A (0.99 ± 0.06, n = 52), 200 nM CLASP1 + 0.5 nM PRC1 + 5 nM Kif4A (0.95 ± 0.15, n = 50), 200 nM + 0.5 nM PRC1 (0.93 ± 0.18, n = 45); 20 nM CLASP1 + 0.5 nM PRC1 + 5 nM Kif4A (0.20 ± 0.26, n = 39); 0.5 nM PRC1 + 5 nM Kif4A (0.0 ± 0.0, n = 50).

Does CLASP1 activity regulate the lengths of crosslinked microtubules in the presence of Kif4A? To answer this question, we lowered the Kif4A concentration to 5 nM where microtubule dynamics reinitiate, and measured the maximum microtubule overlap lengths at varying CLASP1-GFP concentrations. We found that increasing CLASP1-GFP from 0 to 200 nM caused a 7-fold increase in the maximum overlap length (Fig. 4b). Consistent with this, over 90% of all catastrophes are rescued with 5 nM Kif4A and 200 nM CLASP1-GFP (Fig. 4c, Extended data Fig. 6b). Together these data indicate that at low Kif4A (<5 nM in our assay), CLASP1 activity contributes to elongation of microtubule overlaps.

We next examined the dynamics of single microtubules present in the same field of view as the crosslinked bundles. We find that increasing Kif4A concentration systematically from 0-125 nM in the presence of 200 nM CLASP1-GFP reduces the duration-weighted growth rate and maximum microtubule length 2-3-fold, but dynamics is not suppressed (Figs. 4d-4e). Consistent with this, the majority of catastrophe events are rescued under all conditions (Fig. 4f). Further, the trends observed in these quantities were consistent with the dynamics of single microtubules recorded in the absence of PRC1 (Figs. 1c-1d, 1f).

In summary, Kif4A activity is dominant on microtubule overlaps resulting in the formation of stable overlaps under the same conditions where single microtubules continue to grow.

### Why does Kif4A dominate over CLASP1 on crosslinked microtubules?

We performed TIRF assays to simultaneously visualize the localization of CLASP1-GFP and either Alexa 647-labeled Kif4A or Alexa 647-labeled PRC1, starting from the time point of initial encounter between two microtubules to the formation of a stable crosslinked bundle. In experiments with 200 nM CLASP1-GFP, 50 nM Kif4A and 0.5 nM Alexa 647-labeled PRC1, PRC1 was distributed uniformly and enriched rapidly along overlaps (Extended data figs. 7a-7b). Next, the same experiment was performed using 200 nM CLASP1-GFP, 0.5 nM unlabeled PRC1 and either 5 nM or 50 nM Alexa 647-labeled Kif4A. Following the initial encounter between two growing microtubules, both CLASP1 and Kif4A were enriched on the overlap relative to single microtubules (Extended data Fig. 7c-7f and Supplementary video 3). Kif4A localizes to microtubule ends and we observed relative sliding as reported previously^43,44^. CLASP1 is uniformly distributed along the entire overlap, indicating that exclusion of CLASP1 from microtubule ends by Kif4A does not contribute to differential regulation of crosslinked and single microtubules (Extended data fig. 7b, 7d, 7f).

To quantitatively compare the enrichment of CLASP1 and Kif4A on bundles relative to single microtubules, we performed the dynamic bundle assays with GFP-tagged proteins. In assays performed with 0.5 nM PRC1 and 200 nM CLASP1-GFP, CLASP1 showed a 3-fold enrichment on overlaps (Extended data Fig. 8a). In contrast, with 0.5 nM PRC1 and 10 nM Kif4A-GFP, the enrichment of Kif4A was 14-fold on crosslinked microtubules compared to single microtubules (Extended data Fig. 8b).

What leads to these differences in enrichment? The dominant effects of Kif4A could arise from (i) stronger PRC1-Kif4A binding affinity compared to PRC1-CLASP1, or (ii) competition between Kif4A and CLASP1 for PRC1-binding. To investigate the first possibility, we performed BioLayer Interferometry (BLI) assays. We identified the main PRC1-binding region on CLASP1 as the SR-rich region between residues 654-805. The construct CLASP1(654-1471)-GFP bound to full-length PRC1 with a binding K_D_ of 1.19 ± 0.2 μM (Fig. 5a). Under the same conditions, KIf4A-GFP displayed strikingly tighter binding to PRC1 with a K_D_ of 12.47± 2.1 nM (Fig. 5b).

**Figure 5:**
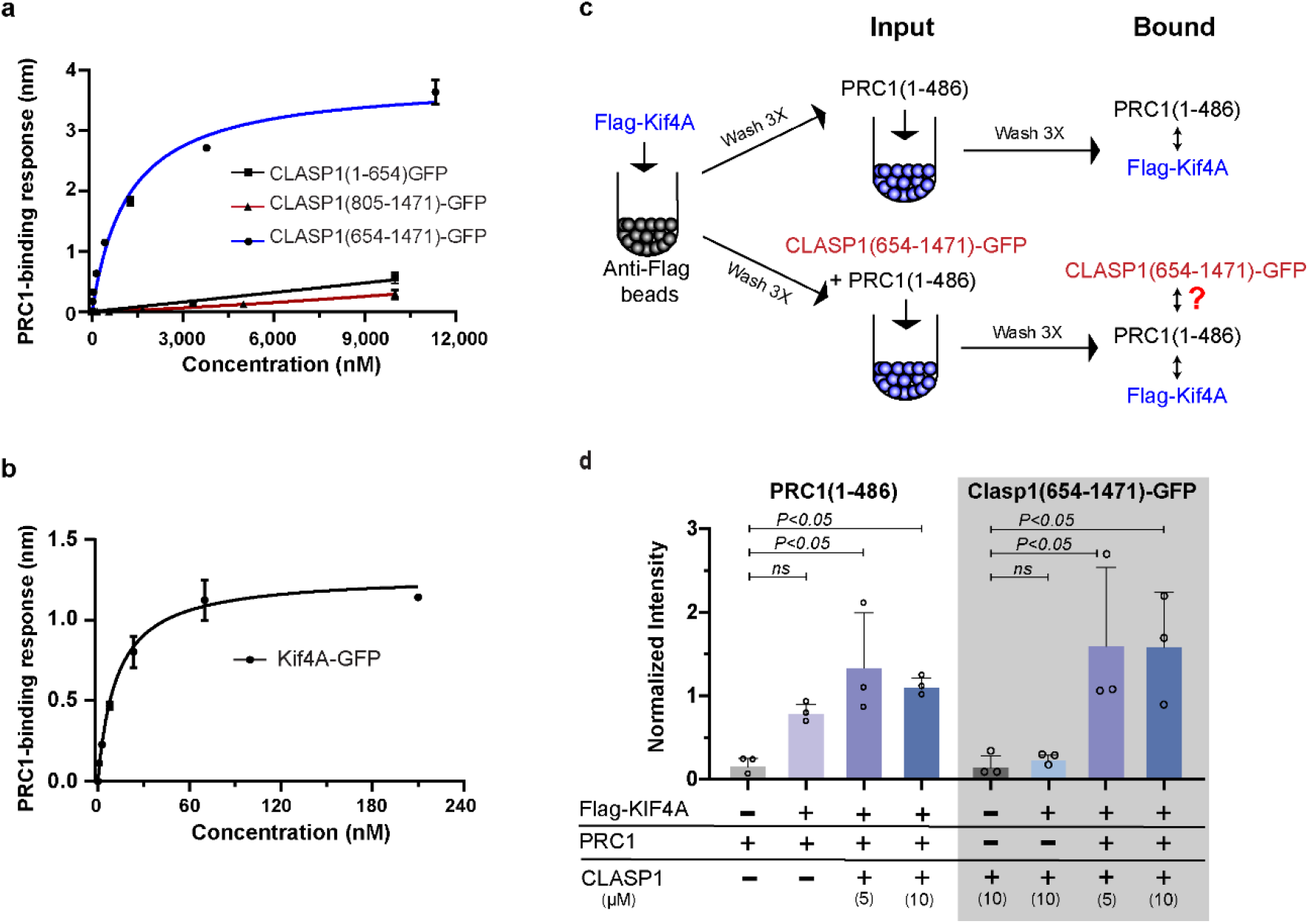
The stronger recruitment of Kif4A to cross-linked overlaps relative to CLASP1 arises from differences in the PRC1-binding affinities of Kif4A and CLASP1. Also see Extended data Figs. 7-10 and Supplementary Video 3 **a.** BLI assay to quantify the binding affinity of CLASP1 constructs to PRC1. Error bars represent standard error of mean. Data collected from 3 independent experiments were fit to a Hill equation. K_D_ : CLASP1(1-654)-GFP > 10 μM; CLASP1(805-1471)-GFP > 10 μM and CLASP1(654-1471)-GFP: 1.19 ± 0.24 μM (R^2^ of fit = 0.98). **b.** BLI assay to quantify the binding affinity of Kif4A-GFP to PRC1. Error bars represent standard error of mean. Data collected from 3 independent experiments were fit to a Hill equation. K_D_ : 12.47± 2.09 nM (R^2^ of fit = 0.98). **c.** Schematic of pull-down assay used to test for competition between Kif4A and CLASP1 for PRC1-binding. Anti-Flag antibody coated magnetic beads were incubated first with Flag-Kif4A, washed and then incubated with either PRC1(1-486) or a combination of PRC1(1-486) + CLASP1(654-1471)-GFP. PRC1(1-486) by itself can bind to Kif4A. For CLASP1(654-1471)-GFP to appear in the Kif4A-bound protein fraction, PRC1(1-486) needs to be able to bind to Kif4A and CLASP1 simultaneously in solution. **d.** Scatter plot with bar graph showing mean of normalized intensities of protein bands of PRC1(1-486) (*left side*) and CLASP1(654-1471)-GFP (*right side, gray box*) from SDS-PAGE gels of pull-down assay described in (c). Error bars indicate standard deviation of intensities from 3 independent experiments. Normalized intensities and standard deviation of PRC1(1-486) bands in sample containing PRC1(1-486): 0.18 ± 0.10; FLAG-Kif4A + PRC1(1-486): 0.80 ± 0.12; FLAG-Kif4A + PRC1(1-486) + 5 μM CLASP1(654-1471)-GFP: 1.36 ± 0.67; FLAG-Kif4A + PRC1(1-486) + 10 μM CLASP1(654-1471)-GFP: 1.12 ± 0.12. P-values for PRC1(1-486) compared to FLAG-Kif4A + PRC1(1-486) + 5 μM and 10 μM CLASP1(654-1471)-GFP are 0.0079 and 0.0252, in an ordinary one-way ANOVA test with Dunnett correction for multiple comparisons. (*Note: the intensity of the PRC1(1-486) band in Flag-Kif4A sample was >3x higher than in sample without Flag-Kif4A in every repeat of 3 independent experiments. The statistical significance in the difference between these samples is difficult to assess due to the high variation in the SDS-gel background relative to the low signal in the control lane without Flag-Kif4A).* Normalized intensities of CLASP1(654-1471)-GFP in sample containing CLASP1(654-1471)-GFP: 0.16 ± 0.15; FLAG-Kif4A + CLASP1(654-1471)-GFP: 0.25 ± 0.07; FLAG-Kif4A- + PRC1(1-486) + 5 μM CLASP1(654-1471)-GFP: 1.62 ± 0.95; FLAG-Kif4A + PRC1(1-486) + 10 μM CLASP1(654-1471)-GFP: 1.60 ± 0.66. P values for CLASP1(654-1471)-GFP compared to FLAG-Kif4A + PRC1(1-486) + 5 μM and 10 μM CLASP1(654-1471)-GFP are 0.0386 and 0.0405 in an ordinary one-way ANOVA test with Dunnett correction for multiple comparisons. P not significant for CLASP1(654-1471)-GFP compared to FLAG-Kif4A + CLASP1(654-1471)-GFP.

To test for competition between Kif4A and CLASP1 for PRC1-binding, we first identified that the CLASP1-binding region on PRC1 lies within residues 1-486 of PRC1 (Extended data Fig. 8c). The construct PRC1(1-486) binds Kif4A with the same affinity as full-length PRC1^34^. We tested if PRC1(1-486) could simultaneously bind to Kif4A and CLASP1 in solution using a pull-down assay, where Kif4A was immobilized on magnetic beads and incubated with either PRC1, CLASP1 or a mixture of the two (Fig. 5c). An analysis of the SDS-PAGE gel of bound proteins showed that CLASP1 (654-1471)-GFP did not bind to Kif4A, but appeared in the Kif4A-bound fraction when it was incubated along with PRC1(1-486) (Fig. 5d, extended data Fig. 8d). This indicates that PRC1 can bind to both KIF4A and CLASP1 simultaneously in solution.

Together these results suggest that higher enrichment of Kif4A compared to CLASP1 on PRC1-crosslinked overlaps does not arise from competition between Kif4A and CLASP1 for PRC1-binding, but stems from the difference in their PRC1-binding affinities.

### Is differential regulation altered in the presence of EB protein?

EB (End-Binding) proteins are known to localize CLASP to tips of dynamic microtubules^36,38^. Therefore, can the presence of EB enrich CLASP1 at tips of crosslinked microtubules so that bundles start elongating, and the differential regulation imparted by the PRC1-Kif4A-CLASP1 module is lost? To answer this question, we systematically included Alexa 647-labeled EB3 in microscopy assays, in combination with the other protein components.

We first examined whether the two microtubule-end localizing proteins EB3 and Kif4A can simultaneously be present on microtubule tips, in a dynamic bundle assay. Kymographs of single microtubules showed that while 50 nM Alexa 647-labeled EB3 alone tracked growing tips, the addition of 5 nM Kif4A resulted in either EB3 or Kif4A being present at microtubule tips, but not both proteins simultaneously. The addition of 50 nm Kif4A almost completely abolished EB3 tip-tracking on both single and crosslinked microtubules, with only Kif4A occupying microtubule ends (Extended data Fig. 9a-c).

The SR-rich domain of CLASP1 contains the PRC1-binding region as well as the EB-binding SxIP motif (Fig. 5a and Extended data fig. 1a). Therefore, we asked if CLASP1 localized with PRC1 at overlaps or at microtubule tips with EB3 when both proteins are present. In dynamic bundle assays with 200 nM CLASP1-GFP, 50 nM Alexa 647-labeled EB3 and 0.5 nM PRC1, we found that CLASP1 is enriched at overlaps. Notably, EB3 did not recruit CLASP1 to the tips of growing microtubules (Extended data Fig. 10a). To understand why, we compared CLASP1-PRC1 and CLASP1-EB3 binding. The K_D_ for CLASP1-PRC1 binding is 1.19 ± 0.2 μM (Fig. 5a). In the absence of salt, the SxIP-motif containing CLASP1(654-1471) protein bound to EB1 with a K_D_ of 1.1 ± 0.3 μM^45^. The binding response of CLASP1(654-1471)-GFP to SNAP-EB3 (EB3 with SNAP-tag) decreased systematically as salt concentration was increased from 10 −50 mM KCl, showing that at 50 mM KCl, binding affinity of CLASP-PRC1 interaction is higher than for CLASP1-EB3 (Extended data Fig. 10b). The recruitment of CLASP1 by PRC1 would be further favored due to the high local concentration of PRC1 at overlaps where it binds cooperatively. Together these results suggested that the addition of EB3 would not alter differential regulation of single and crosslinked microtubules in the CLASP1-PRC1-Kif4A system. Subsequent dynamic bundle assays with CLASP1, Kif4A, PRC1 and EB3 confirmed that this is indeed the case (Extended data Fig. 10c)

## DISCUSSION

It is well established that the length of a microtubule or the overlap between two microtubules can be tuned by the collective activity of MAPs with opposite functions, such as tubulin polymerases and depolymerases, or inward-outward sliding by plus- and minus-end directed motors^46^. This study illustrates that in addition to these canonical mechanisms of length regulation that operate on a single microtubule or one type of array, the collective activity of MAPs on an ensemble of microtubule arrays can differentially regulate the dynamics and size of its constituent sub-populations.

We find that a minimal protein module comprised of three MAPs: the antiparallel crosslinker PRC1, a kinesin that suppresses microtubule dynamics Kif4A, and a rescue factor CLASP1, can establish a system where single microtubules predominantly elongate while tubulin polymerization is completely suppressed in crosslinked bundles under identical reaction conditions (Fig 6a). In contrast to a simple model where differential regulation arises by restriction of CLASP1 localization to single microtubules and Kif4A to crosslinked overlaps, we find that all proteins are present on both microtubule populations. The length regulation of both single and crosslinked microtubules is neither a simple function of the Kif4A/CLASP1 ratio in solution, nor directly related to the lattice-bound concentration of either protein (Figs. 1d,1e,4b and 4e). Instead, it is determined by a combination of multiple parameters such as the geometry of microtubule arrays, microtubule- and PRC1-binding affinities and intrinsic activity of regulators on dynamic microtubules.

**Figure 6:**
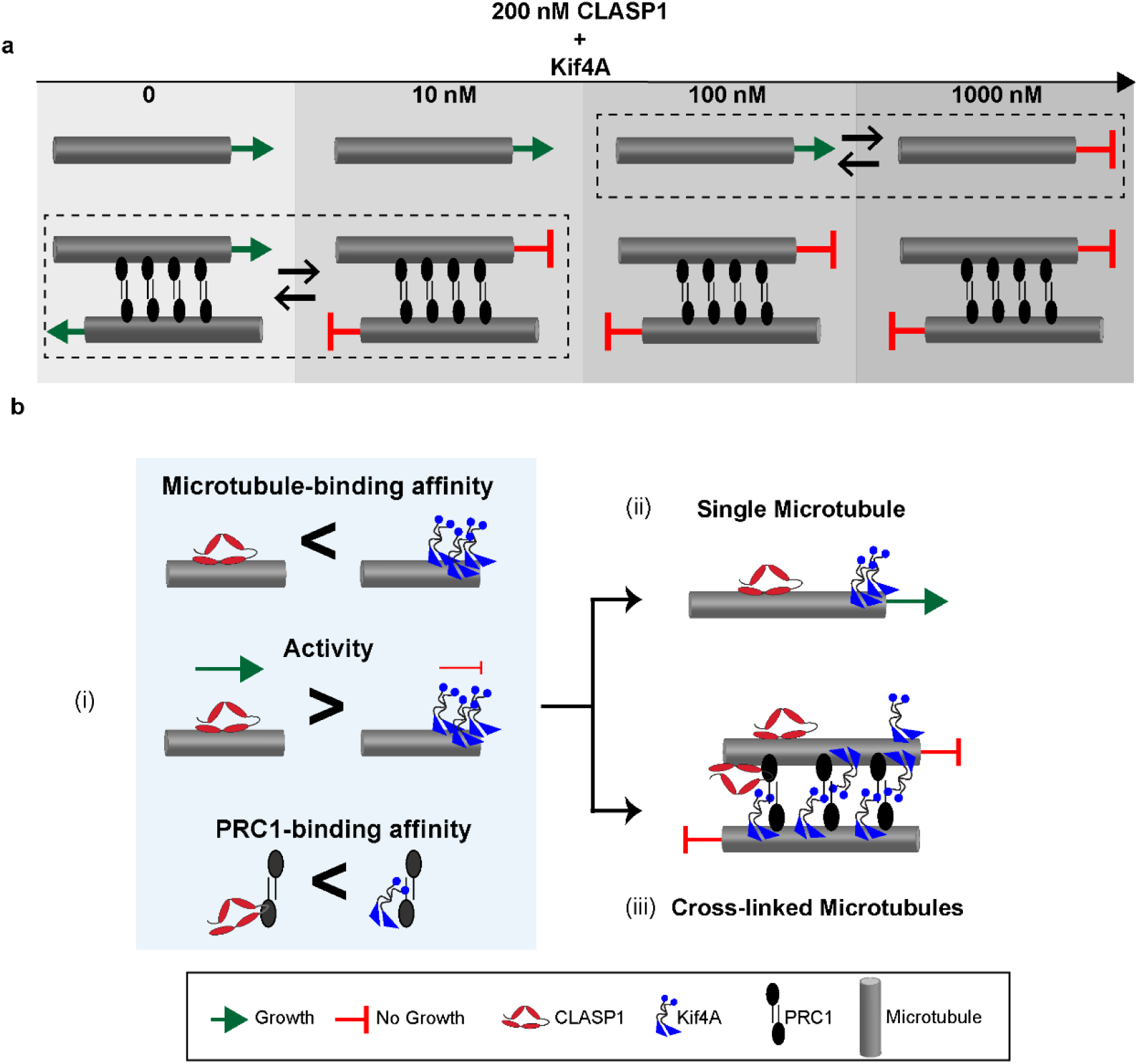
An inverse microtubule affinity-activity relationship along with differences in PRC1-binding affinity between CLASP1 and Kif4A underlie differential regulation of single and cross-linked microtubules by the PRC1-CLASP1-Kif4A protein module. **a.** The dynamics of single microtubules (top) and cross-linked microtubules (bottom) are differentially regulated by the PRC1-CLASP1-Kif4A protein module. Box with dotted lines indicates regime where dynamics switch between continuous growth (→) and no elongation (┤). In the presence of constant CLASP1 concentration, the switch for each microtubule populations occurs at strikingly different Kif4A concentrations. **b.** (i) Differences in three properties of CLASP1 compared to Kif4A underlie this differentiation: Lower intrinsic microtubule-binding affinity of CLASP1 relative to Kif4A, higher intrinsic activity of CLASP1 relative to Kif4A, and higher PRC1-binding affinity of Kif4A compared to CLASP1. As a result, the high activity of CLASP1 as a rescue factor overcomes Kif4A activity on single microtubules, despite lower CLASP1 microtubule affinity. However, on cross-linked microtubules, the enrichment of Kif4A through its stronger PRC1-binding and microtubule-binding helps it overcome CLASP1 activity. The low affinity of CLASP1 for microtubules permits Kif4A activity to suppress the growth of crosslinked microtubules.

How are single and crosslinked microtubules differentially regulated by the PRC1-Kif4A-CLASP1 module? We find that on single microtubules, CLASP1 activity dominates to promote rescue and thereby microtubule elongation even at a 5-fold lower concentration than Kif4A (Fig. 1). In this regime, microtubule-binding of CLASP1 is at least 4-9 fold weaker compared to Kif4A (Fig. 1e), which could arise from the transient nature of CLASP binding to the lattice^36,38,40^. We propose that similar to its isoform CLASP2, a low number of CLASP1 molecules bound along the microtubule lattice are sufficient to initiate a rescue, whereas high concentrations of Kif4A at the tips is needed to suppress microtubule dynamics^36^. This raises the question: how do low nanomolar concentrations of Kif4A overcome CLASP1 activity on crosslinked microtubules? The change in dynamics arises primarily from the 10-28 fold greater accumulation of PRC1^35,47^ and the subsequent 14-fold enrichment of Kif4A that we observe at overlaps compared to single microtubules (Extended data Fig.8b). In comparison, CLASP1 is enriched only 3-fold at overlaps over single microtubules, due to lower PRC1-binding affinity relative to Kif4A. In combination with the lower microtubule-affinity of CLASP1, growth suppression by Kif4A dominates on crosslinked microtubules. The concentration of PRC1 could serve as an additional knob to fine-tune the dynamics in this system, by altering the switch points shown in Fig.6a.

These studies show that in addition to differences in PRC1-binding affinity, an inverse relationship between intrinsic properties of Kif4A and CLASP1, i.e., lower activity-higher microtubule affinity of Kif4A and higher activity-lower microtubule affinity of CLASP1, is essential for differential regulation (Fig. 6b). While the contribution of PRC1-binding affinity to differential regulation is intuitive, the importance of the inverse relationship between the intrinsic microtubule-binding affinity and activity of the two regulators can be better understood by considering possible outcomes in its absence. For example, in addition to lower PRC1-binding affinity, if CLASP1 had high microtubule binding affinity, then its high activity would result in the elongation of bundles as well as single microtubules. Similarly, if Kif4A had both high activity and high microtubule binding affinity relative to CLASP1, both single and crosslinked microtubules would elongate less.

In contrast to a mechanism involving spatial segregation of proteins into different arrays, the concurrent activity of both Kif4A and CLASP1 on single and crosslinked microtubules as described here, has several advantages. First, tight spatial partitioning is difficult to achieve in dynamic systems where proteins are characterized by fast turnover on microtubules, as is the case for PRC1, CLASP1 and Kif4A at the cell center^14,25,48^. Second, this system permits the modulation of stability and length of single and crosslinked microtubules by both regulators. Third, the presence of both proteins allows individual microtubule arrays to independently switch between dynamic states. This property would be advantageous in re-initiating microtubule growth in case of damage to either array. Finally, the simultaneous presence of all three proteins imparts remarkable robustness to the system, as exemplified by experiments with EB proteins, where we find that the protein-protein interactions that characterize the CLASP1-PRC1-Kif4A module are not perturbed by addition of EB (Extended data figs. 9 & 10). Thus, the characteristic features of the PRC1-Kif4A-CLASP1 module presents a highly robust, yet versatile and tunable system.

How might such a module, which consists of regulators with opposite effects on microtubule dynamicity and lengths, promote microtubule organization at the cell center during anaphase? Features of the anaphase spindle differ widely across cell types, with the number of inter-polar microtubules varying from fewer than 10 to many 100s, and with spindle elongation lengths ranging from 1 to 10 μm^49^. In the face of such diversity, the presence of both the growth promoter and suppressor on single and crosslinked microtubules provides a flexible strategy whereby the dominant regulator could change depending on the spindle type and the need of the system. This is seen in systems such as *Xenopus* egg extracts and *Drosophila* spermatocytes at anaphase where CLASP homologs increase the stability of all spindle microtubules, including bundles^24,50^. In fission and budding yeast, the spindle localization of the growth suppressors of the Kinesin-8 family is independent of the cross-linker Ase1p^22,51^. This may reflect an adaptation suited to the large elongation of its spindle, which is chiefly mediated by the activity of rescue factors and polymerases such as CLASP and XMAP215 homologs on crosslinked microtubules^23,51^. In yet other systems, the nucleation and elongation of non-centrosomal microtubules at anaphase onset contribute significantly to spindle assembly. In these systems, CLASP activity on single microtubules is essential for initial microtubule assembly^32^. Together, the three-protein module provides an adaptable, versatile platform which can promote the nucleation and elongation of new microtubules, and tune the lengths and stability of single and crosslinked arrays separately, through differential regulation of their dynamics.

On a broader note, the “inverse activity-microtubule affinity relationship” that underlies differential regulation does not rely on features specific to the anaphase spindle, and these design principles can be extended to other cytoskeletal structures across the cellular cytoplasm.

## METHODS

### Protein expression and purification

Full-length human CLASP1 protein (GenBank: BC112940.1) and all deletion constructs (CLASP1(1-654), CLASP1(654-1471) and CLASP1(805-1471)) were cloned into a modified pFastBac expression vector (Thermo Fischer Scientific) that contained a PreScission Protease cleavable N-terminal Twin-Strep-Tag and a C-terminal GFP followed by a Tobacco etch Virus (TEV) cleavable 6x-His tag. Proteins were expressed in the Sf9 insect cell line using the Bac-to-Bac^®^ Baculovirus Expression System (Thermo Fischer Scientific), and cells were grown in HyClone™ CCM3 SFM (GE Life Sciences). Pellets of all deletion constructs were lysed by a short sonication in buffer A (50 mM Phosphate buffer pH 8.0, 300 mM NaCl, 10 % glycerol, and 30 mM imidazole) supplemented with 0.15 % tween, 0.5 % Igepal 630, 2 mM TCEP, 1 mM PMSF, 0.5 mM Benzamidine Hydrochloride, 75 U Benzonase and protease inhibitor cocktail (Thermo Scientific). The lysate was clarified by ultracentrifugation (70,000 × g, 30 min) and supernatant was incubated with Ni-NTA resin (Qiagen) for 1 hour. The resin was washed with buffer A supplemented with 0.15 % tween and 0.5 mM TCEP and bound protein was eluted with 50 mM Phosphate buffer pH 8.0, 300 mM NaCl, 5 % glycerol and 400 mM imidazole. Peak protein fractions were pooled and incubated with TEV and PreScission protease (30:1 w/w of protein:protease) overnight at 4 °C. The proteins were further purified by size exclusion chromatography (SEC; HiLoad 16/600 Superdex 200 pg, GE Healthcare) in 50 mM phosphate pH 8.0, 300 mM NaCl, 5 % glycerol and 5 mM β-mercaptoethanol, and flash-frozen in liquid nitrogen. Full-length CLASP1-GFP was purified similar to the protocol described above, with the following modifications: Lysis buffer contained 50 mM Phosphate buffer pH 7.5, 500 mM NaCl, 10 % glycerol, 30 mM imidazole, 100 mM arginine, supplemented with 0.15 % tween, 0.5 % Igepal 630, 2 mM TCEP, 1 mM PMSF, 75 U Benzonase and protease inhibitor cocktail (Thermo Scientific). Following a short sonication and ultracentrifugation, the clarified supernatant was passed through a Ni-NTA superflow cartridge (Qiagen). After elution, fractions containing pure protein were pooled and diluted with elution buffer, such that protein concentration did not exceed 1 mg/ml. The 6x-His tag was cleaved with TEV protease in the presence of 100 mM arginine, 2 mM β-mercaptoethanol, 5 mM EDTA and 0.1 % tween. The cleaved protein was further purified through SEC in a Superose 6 increase 10/300 GL column (GE Healthcare), in buffer containing 50 mM phosphate pH 7.5, 500 mM NaCl and 10% glycerol. Kif4A-GFP, Kif4A^34^ and PRC1^47^ were purified as previously described. Kif4A-CLIP and SNAP-PRC1 were purified similar to their respective full-length proteins.

Full-length human EB3 gene was expressed in Rosetta (DE3) cells from a pST50Tr-HISDHFR plasmid, with an N-terminal His-tag followed by a TEV cleavage site and the SNAP protein (Plasmid from Julie Welburn’s lab, University of Edinburgh, Scotland). Protein expression was induced by the addition of 0.1 mM IPTG, and cells were grown overnight at 18°C and centrifuged. Frozen cell pellets were thawed and lysed by sonication in PBS buffer at pH 8.0, supplemented with 30 mM Imidazole, 0.15 % tween, 0.5 % Igepal 630, 2 mM TCEP, 1 mM PMSF, 0.5 mM Benzamidine Hydrochloride, 75 U Benzonase and protease inhibitor cocktail (Thermo Scientific). Lysate was clarified through ultracentrifugation at 80,000xg for 40 minutes, and the supernatant was incubated with Ni-NTA resin for 1 hour. The resin was washed with lysis buffer and bound-protein was eluted with lysis-buffer supplemented with 400 mM Imidazole. Subsequently, cleavage of the purification tag and further purification by SEC through a Superdex 200 10/300 GL column were performed as described for CLASP deletion constructs.

Tubulin was purified^52^ or purchased from either PurSolutions, LLC or Cytoskeleton, Inc.. Tubulin was labeled in 1:10 proportions with biotin or X-rhodamine according to published protocols or mixed with prelabeled tubulin purchased from Cytoskeleton, Inc.

### Fluorescent labeling of proteins

CLIP and SNAP tagged proteins were labeled using CLIP Surface-647 and SNAP Surface-647 dyes (New England Bio Labs), respectively. Purified protein at a concentration of 2 mg/ mL was incubated with 10 mM β-mercaptoethanol and 4x molar excess of dye at room temperature for 5 min, and then on ice overnight. Unlabeled dye was removed through dialysis in a Mini Dialysis Kit, 8 kDa cut-off (Cytiva) for 4 hours at 4°C, in buffer containing 50 mM phosphate pH 8.0, 300 mM NaCl, 5% glycerol, 30% sucrose, 10 mM β-mercaptoethanol, followed by multiple rounds of concentration and dilution through an Amicon Ultra 0.5 mL concentrator (Sigma-Aldrich). Labeling efficiency was calculated by measuring absorbance at 280 nm and 650 nm (A_280_ and A_650_) and using extinction coefficients of protein and dye (ε_280_ and ε_650_), as

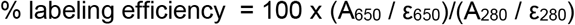

The labeling efficiency for the proteins used in TIRF assays were Alexa 647-labeled Kif4A: 47%; Alexa 647-labeled PRC1: 17% and Alexa 647-labeled EB3: 45%.

### Solution light scattering studies

For SEC-MALS studies, between 600-900 μg of each CLASP1 protein was injected into a Superose 6 column (GE Healthcare) in a buffer containing 50 mM phosphate pH 7.6, 200 mM NaCl, 5% Glycerol and 1 mM DTT. The column was connected to High Performance Liquid Chromatography System (HPLC), Agilent 1200, (Agilent Technologies, Wilmington, DE) equipped with an autosampler. The elution from SEC was monitored by a photodiode array (PDA) UV/VIS detector (Agilent Technologies, Wilmington, DE), differential refractometer (OPTI-Lab rEx Wyatt Corp., Santa Barbara, CA), static and dynamic, multiangle laser light scattering (LS) detector (HELEOS II with QELS capability, Wyatt Corp., Santa Barbara, CA). Two software packages were used for data collection and analysis: the Chemstation software (Agilent Technologies, Wilmington, DE) controlled the HPLC operation and data collection from the multi-wavelength UV/VIS detector, while the ASTRA software (Wyatt Corp., Santa Barbara, CA) collected data from the refractive index detector, the light scattering detectors, and recorded the UV trace at 280 nm sent from the PDA detector. The weight average molecular masses, Mw, were determined across the entire elution profile in the intervals of 1 sec from static LS measurement using ASTRA software as previously described^53^. Hydrodynamic radii, Rh, were measured from an “on-line” dynamic LS measurement every 2 sec. The dynamic light scattering signal was analyzed by the method of cumulants^54^.

The SEC-LS/UV/RI instrumentation was supported by NIH Award Number 1S10RR023748-01. The content of this paper is solely the responsibility of the authors and does not necessarily represent the official views of the National Institutes of Health.

### Total Internal Reflection Fluorescence Microscopy Assays

Microscope chambers were constructed using a 24 × 60 mm PEG-Biotin coated glass slide and 18 × 18 mm PEG coated glass slide separated by double-sided tape to create two channels for exchange of solutions. Standard assay buffer was 1 × BRB80 (80 mM K-PIPES at pH 6.8, 2 mM MgCl_2_ and 1 mM EGTA at pH 6.8), 50 mM KCl, 1 mM ATP, 1 mM GTP, 0.1 % methylcellulose and 3 % sucrose. Imaging was carried out in the presence of an antifade reagent mix (40 mg/ml Glucose Oxidase, 35 mg/ml Catalase, 25 mM Glucose, 0.5% β-mercaptoethanol and 10 mM Trolox reagent). Images were acquired using NIS-Elements (Nikon) and analyzed using ImageJ. Data were analyzed from experiments performed on three independent days, unless specified otherwise.

#### Dynamic Microtubule Assay of Single Microtubules

Experiments with dynamic microtubules were carried out as described in Jiang *et al*., 2019^45^. X-rhodamine (1:10 labeled to unlabeled) and biotin (1:10 labeled to unlabeled) labeled microtubules were polymerized in the presence of GMPCPP, a non-hydrolysable GTP-analogue, and immobilized on a neutravidin coated glass coverslip. Coverslips were briefly incubated with casein to block non-specific surface binding. A low concentration of CLASP1-GFP (1 nM) was added and accumulation of protein preferentially to the plus-end of seed was used as an indicator of microtubule polarity. Subsequently, 16 μM tubulin (1 X-rhodamine-labelled tubulin:10 unlabeled tubulin) in assay buffer and antifade reagent along with CLASP1 and Kif4A protein and ATP were added. Images were recorded every 10 seconds for 20 minutes. In all microtubule experiments, concentrations of monomeric CLASP1 and dimeric Kif4A are reported, unless otherwise specified.

#### Dynamic Microtubule Assay of Microtubule Bundles (“Dynamic bundle assay”)

HiLyte647 (1:10 labeled to unlabeled) and biotin (1:10 labeled to unlabeled) labeled microtubules were polymerized in the presence of GMPCPP, a non-hydrolysable GTP-analogue, and immobilized on a neutravidin coated glass coverslip. Coverslips were briefly incubated with casein to block non-specific surface binding. Microtubule bundles were formed through the addition of 5 nM PRC1 and GMPCPP-polymerized X-rhodamine microtubules. CLASP1, Kif4A, PRC1 and tubulin were subsequently added in the presence of antifade reagents. Images were recorded every 10 seconds for 20 minutes. For experiments performed to quantify the enrichment of Kif4A-GFP on bundles in the absence of CLASP1, data from 2 sets of experiments from two independent days of experiments were analyzed. In all microtubule experiments, concentrations of monomeric CLASP1 and dimeric Kif4A and PRC1 are reported, unless otherwise specified. For experiments performed with EB3, dynamic bundle assays were performed as described above, with the only difference being the use of X-rhodamine-labeled GMPCPP microtubule seeds. For images shown in Extended data fig. 6c, kymographs from 45 bundles from 3 independent days of experiments were visually examined and a representative kymograph was chosen. Similarly, for Extended data figs. 7a and 7c, 73 and 67 kymographs were respectively examined.

#### Simultaneous visualization of all proteins on bundles

A high density of X-rhodamine labeled, GMPCPP microtubule seeds were immobilized on the coverslip as described above. Microtubule polymerization was initiated by the addition of X-rhodamine labeled tubulin, 50 mM KCl, 1 mM ATP, GTP, CLASP1-GFP and a combination of either Alexa 647-labeled Kif4A-CLIP and unlabeled PRC1, or Alexa 647-labeled SNAP-PRC1 and unlabeled Kif4A. Data were collected from 2 independent experiments for CLASP1-GFP + Alexa 647-labeled PRC1 + Kif4A, and PRC1 localization and enrichment were found to be consistent with published reports. Data were collected from 3 independent experiments for CLASP1-GFP + Alexa 647-labeled Kif4A + PRC1. All videos from 3 independent experiments were visually analyzed and representative bundles where 2 growing tips encounter each other to form a crosslinked overlap were chosen for the montages and line-scans shown in Extended data fig. 7. Time point t=0 was set at the first frame where microtubules encountered each other, as judged from the intensity in the X-rhodamine channel. For line-scans, intensities were normalized from 0 (0 intensity) to 100 (maximum intensity recorded over all time points in each individual channel).

#### Single molecule imaging

Microtubule seed immobilization and polymerization were performed as described in the section “Dynamic Microtubule Assay of Single Microtubules”. For single molecule imaging, assays were performed in the absence of KCl. A time-lapse sequence of images was acquired at a rate of 0.3 frames/s using an ANDOR iXon Ultra EMCCD camera. Kif4A concentrations in these experiments refer to the monomer.

#### Kif4A-GFP and CLASP1-GFP binding to single microtubules

HiLyte647 (1:10 labeled to unlabeled) and biotin (1:10 labeled to unlabeled) labeled microtubules were polymerized in the presence of GMPCPP and taxol, and immobilized on a neutravidin coated glass coverslip. Coverslips were briefly incubated with casein to block non-specific surface binding. Kif4A-GFP or CLASP1-GFP were subsequently added, along with standard assay buffer, antifade reagent, ATP and 50 mM KCl. For steady-state analysis of GFP intensity, Kif4A-GFP and CLASP1-GFP were incubated with immobilized microtubules for 5 min before still images were recorded. Data from two independent days of experiments were combined. Data from a third day was collected and analyzed and showed consistently higher intensity values owing to changes in the microscope’s settings. These data were separately analyzed and showed the same trends as data from the other two sets. Concentrations of monomeric Kif4A-GFP were used in these experiments to enable direct comparison between CLASP1-GFP and Kif4A-GFP intensities.

### Quantification and Statistical Analysis

Kymographs from growing microtubules were generated using the multiple kymograph plugin in ImageJ. Quantitative analyses were carried out in GraphPad Prism. “n” numbers in all experiments refer to the unique number of microtubules used for the dataset and standard deviations correspond to deviations from the mean. Ordinary one-way ANOVA tests with Dunnett correction for multiple comparisons were perfomed on GraphPad Prism to determine P-values for statistical significance.

#### Growth Rate Analysis

For each event in a kymograph, growth rates were calculated as:

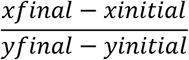

The output was the growth rate in nm/s. These values were used to plot the growth rate of each event and the distribution of growth rates.

#### Distribution plot of growth rates

Individual growth rates from each event on a kymograph were plotted on a distribution plot using GraphPad Prism. Relative frequency values were plotted with a bin width of 3 nm/s.

#### Duration-weighted growth rate

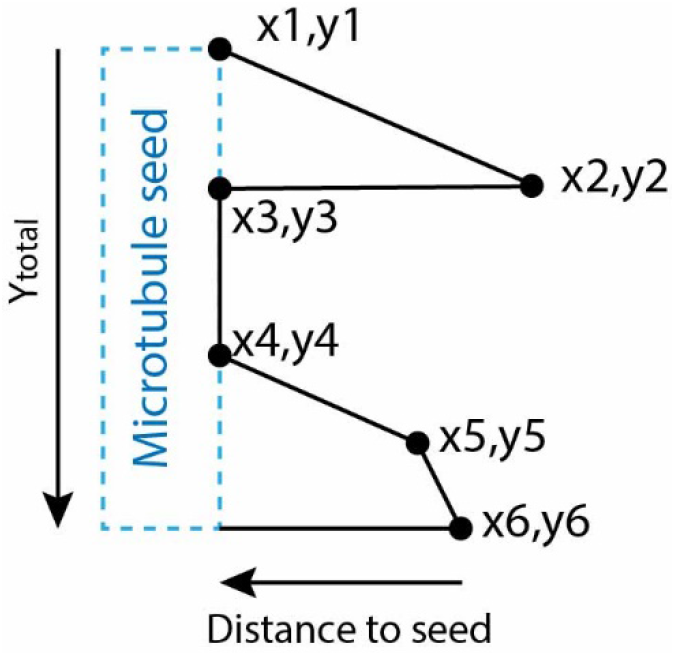

For the illustrative kymograph shown above, the growth rate over each phase is uniform, and depolymerization events are rapid and occur within the time gap between successive images.

The growth rates for each of the 4 phases thus become 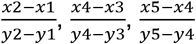 and 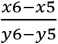.

The average growth rate over the entire kymograph, accounting for the differences in duration of each phase equates to:

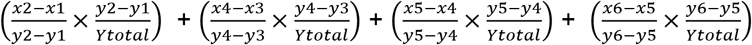

In general, the average growth rate of a microtubule weighted by the duration of each of the ‘n’ phases it goes through is

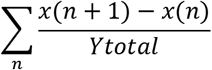

#### Fraction of time microtubule stalled plot

Instantaneous growth rates were recorded and all values <3 nm/s were considered as stalling events. (Pixel size of the Zyla CMOS camera is 65 nm. Frames were recorded every 10 seconds for all images. Therefore at 3 nm/s, there would only be 30 nm growth which is less than 1 pixel). The fraction of time each microtubule stalled was calculated as below:

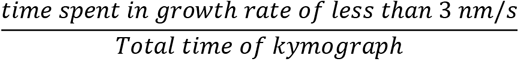

#### Average microtubule length before catastrophe event analysis

Microtubule lengths were measured from the tip of a growing microtubule to the seed for each catastrophe event. Length at the last timepoint was excluded from the calculation as no catastrophe could be recorded. Each individual length from all kymographs analyzed were plotted.

#### Maximum microtubule length analysis

For each kymograph, the maximum length each microtubule reached was recorded. Lengths were measured from the tip of the microtubule to the seed. Length at the last timepoint was included. Each individual value for each kymograph was plotted.

#### Average number of rescues

For each kymograph, the number of rescue events were recorded. Rescue events were defined as catastrophe events which did not depolymerize to the GMPCPP microtubule seed.

#### Rescue as a fraction of total number of catastrophes

For each kymograph, the number of rescue events and catastrophe events were recorded. Rescue events were defined as above and catastrophe events were defined as all depolymerization events. The frequency of rescue events as a fraction of total number of catastrophe events were plotted for each individual kymograph.

#### Intensity analysis

ImageJ was used to assess GFP fluorescence intensities on microtubules. For all average intensity per pixel values recorded, a rectangular area along the microtubule was selected with a width of 5 pixels. Background intensities were also subtracted locally from regions of interest using a box of the same size close to the selected microtubule. Intensities were not analyzed for microtubules found at the edges of the camera’s field of view.

#### Single molecule GFP Intensities

Tip intensity was determined for individual growth events from kymographs using a 7-pixel wide line. The same line was also drawn on a region of the microtubule lattice to determine lattice intensity. Total intensity was calculated as the sum of the tip intensity and lattice intensity. The intensity values were corrected by subtraction of the mean intensity of the background.

#### EB3 tip intensities

The tip intensity for EB3 in the presence of Kif4A was determined over a region of fixed length and 7-pixel width for individual growth events. The intensity values were corrected by subtraction of the mean intensity of the background.

### BioLayer Interferometry (BLI) Assays

BLI experiments were performed in an Octet Red 96 instrument (ForteBio). Full length PRC1 or PRC1(1-486) (the ‘ligand’ protein) was immobilized on an Amine-Reactive Second-Generation (AR2G) sensor at a concentration of 12 μg/ml in sodium acetate buffer pH 5, using the AR2G Reagent kit (ForteBio). The analyte proteins (CLASP1(1-654)-GFP, CLASP1(1-805)-GFP, CLASP1(805-1471)-GFP and Kif4A-GFP) were diluted in binding buffer: 1X BRB80 (80 mM K-PIPES at pH 6.8, 1 mM MgCl_2_ and 1 mM EGTA at pH 7.2) supplemented with 50 mM KCl, 1 mM DTT and 0.1 % tween, at concentrations ranging from 0-10 μM for CLASP proteins and 0-120 nM for Kif4A-GFP. Ligand-bound sensors were first dipped into quenching solution (1 M ethanolamine), and then in pre-blocking solution containing binding buffer with 1 mg/ml α-casein, to reduce non-specific binding of analyte. Following this, they were dipped briefly in binding buffer and then into analyte proteins to record the binding response for association of analyte to ligand, and then into plain binding buffer for dissociation. The assay was repeated sans ligand protein to measure non-specific binding of analyte to the sensor. Data were analyzed using Data Analysis 9.0 software (ForteBio) and double referencing was performed to correct for drift of ligand from sensor and non-specific binding of analyte. Binding responses at 180s in the association step from three independent experiments were plotted against analyte concentrations and data were fit to a Hill equation to determine the binding K_D_.

For EB3-CLASP1 experiments, master stocks of 20 μg/ml SNAP-EB3 in sodium acetate buffer pH 5 (ligand protein), and 1 μM CLASP1(654-1471)-GFP (analyte) in binding buffer without KCl were prepared. Proteins were aliquoted into wells of a 96 plate for BLI experiments, and salt concentrations were changed by the addition of KCl. The assay was performed as described above and ch anges in the binding response of CLASP1with salt were verified from three independent experiments.

### Pull-down assays

Anti-Flag M2 magnetic beads (Sigma Aldrich) were equilibrated in assay buffer containing 1X BRB80 at pH 7.2, 50 mM NaCl, 0.1% CHAPS, 1 μg/ μl of α-casein and 0.5 mM TCEP. Kif4A-CLIP protein with an N-terminal Flag tag, at a concentration of 2.5 μM in assay buffer with 100 μM ATP was immobilized on equilibrated beads placed on ice. The beads were then washed 3 times with an excess of assay buffer and all liquid from the tubes were removed using a magnetic separation rack. Reaction mixtures containing either 5 μM PRC1(1-486), 5 or 10 μM CLASP1(654-1471)-GFP or combinations of the two proteins in assay buffer were incubated with the beads for one hour. To measure non-specific binding of PRC1 and CLASP1, equal volumes of reaction mixtures were incubated with equilibrated beads that were not bound to Flag-Kif4A-CLIP. Following incubation, unbound proteins were collected from the beads and mixed with SDS-gel loading dye. The beads were then washed with assay buffer 3 times and then incubated at 98°C for 3 min in the presence of SDS-gel loading dye to elute bound proteins. Protein samples were run on an NOVEX WW 4-12% TG GEL (Thermo Fisher Scientific), and gels were stained with Coomassie. Intensities of gel bands were quantified by measuring absorbance at 635 nm on an Amersham™ Typhoon™ Biomolecular Imager (Cytiva life sciences). Measurements of gel bands of known concentrations of BSA under the same gel loading, staining and imaging conditions confirmed that the detector showed a linear response in the range of intensities being imaged.

#### Quantitative analysis of gel intensities

In order to account for differences in the volume of beads per reaction and the volume of sample loaded, intensities of all bands were normalized against the intensity of anti-flag antibody present in the same sample. Normalized intensities from three independent pull-down experiments were plotted, and differences between band intensities were tested for statistical significance using an ordinary one-way ANOVA test on GraphPad Prism.

## ACKNOWLEDGEMENTS

This work was supported by a grant from the NIH (1DP2GM126894-01), and funds from the Pew Charitable Trusts and the Smith Family Foundation to R.S. We thank Julie Welburn (University of Edinburgh, Scotland) and Marija Zanic (Vanderbilt University, USA) for the generous gift of EB plasmids, and Bob K. Dass (ForteBio, Sartorius) for help with troubleshooting BLI assays. The SEC-MALS instrumentation was supported by NIH Award Number 1S10RR023748-01 to Ewa Folta-Stogniew (Yale School of Medicine, USA).

