## Extended data fig for "Differential Regulation of Single Microtubules and Bundles by a Three-Protein Module"

### Extended Data Fig. 1

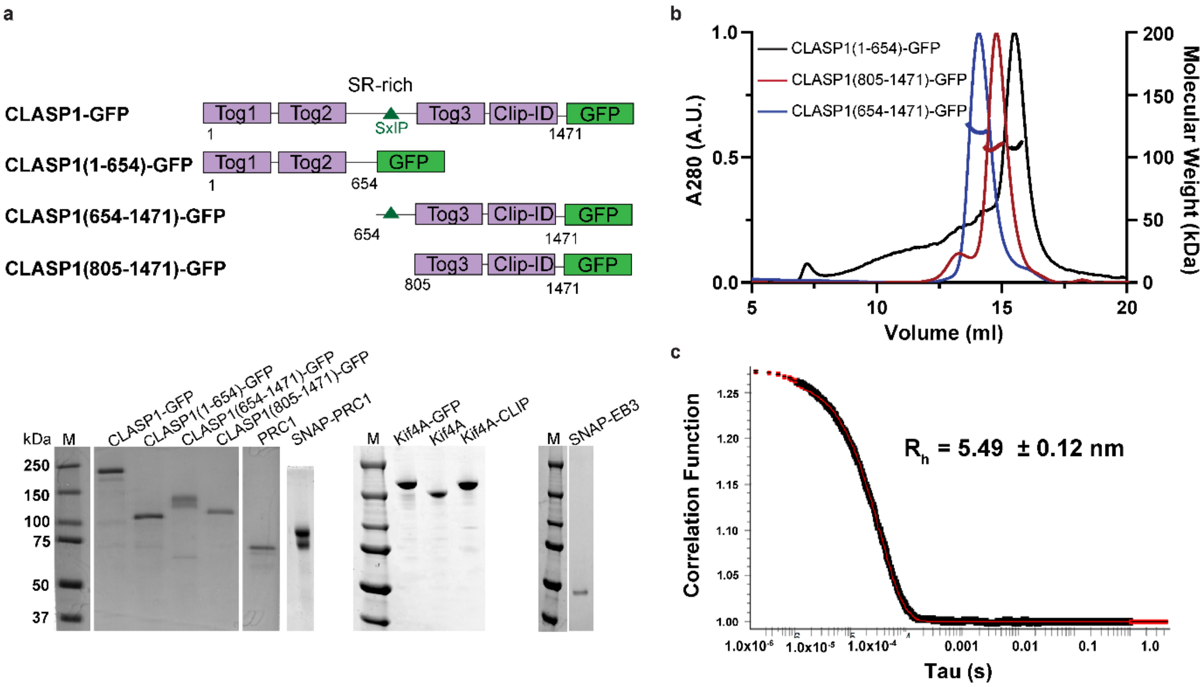


**Extended Data Fig. 1.** *Related to Fig. 1*

**a.** Domain diagram of CLASP1 constructs used in this study (top), along with SDS-PAGE gel of all purified proteins used in this study (bottom). SR-rich: Serine Arginine rich, CLIP-ID: CLIP-Interacting Domain, M: molecular weight marker, with corresponding masses in kDa on the side.

**b.** SEC-MALS profiles showing elution volumes of CLASP1 constructs from a Superose-6 column, and their corresponding molecular weights in solution. *Mean and standard deviation of calculated Molecular weight (expected weight of monomer):* CLASP1(1-654)-GFP (black) 108.8 ± 0.8 kDa (101.5 kDa); CLASP1(805-1471)-GFP (red) 107.6 ± 10.8 kDa (102.2 kDa); CLASP1(654-1471)-GFP (blue):128.7 ± 0.3 kDa (120 kDa).

**c.** Auto-correlation function for apex of eluting peak of CLASP1(805-1471)-GFP shown in (**b**), obtained through dynamic light scattering studies. Calculated hydrodynamic radius, $R_h$= 5.49 ± 0.12 nm.

### Extended Data Fig. 2

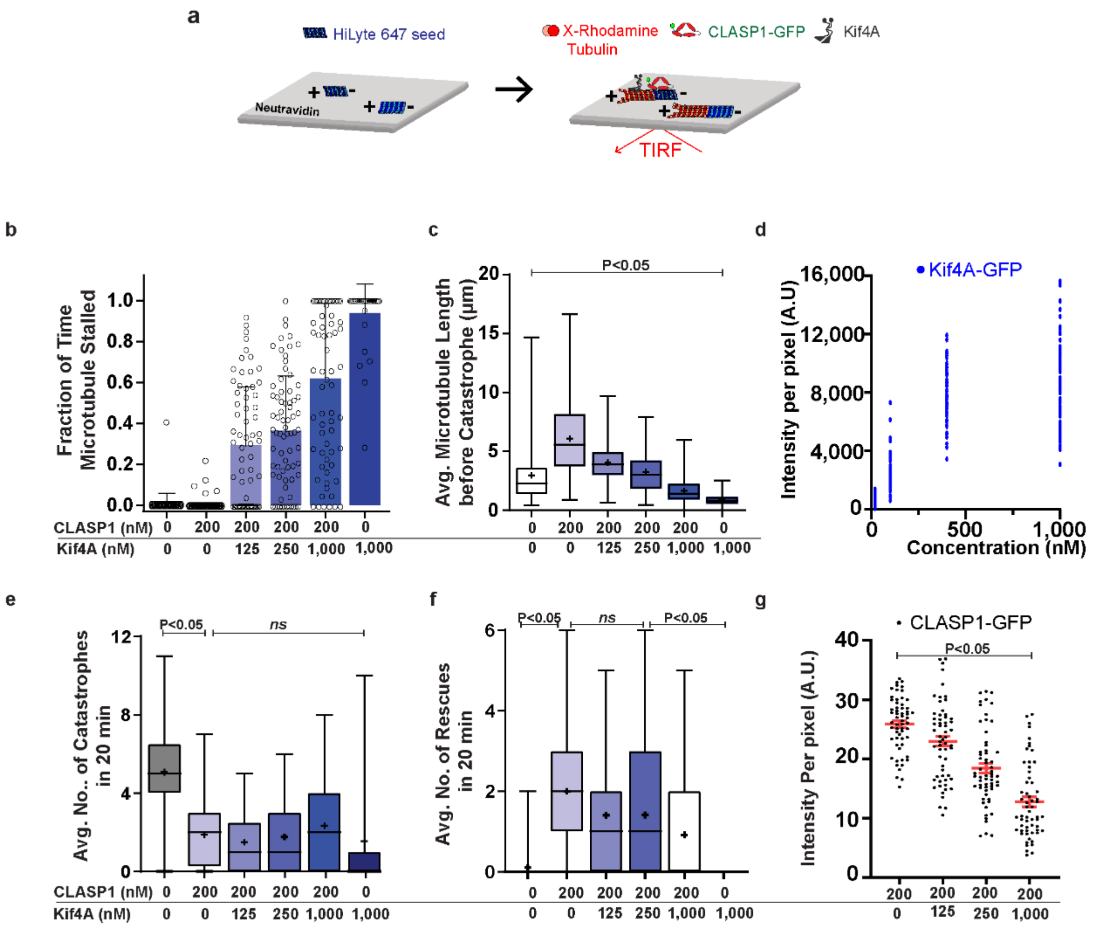


**Extended Data Fig. 2.** *Related to Fig. 1*

n = number of kymographs analyzed from 3 independent experiments, except where indicated.

P values were calculated from an ordinary one-way ANOVA test with Dunnett correction for multiple comparisons.

**a.** Schematic of the dynamic microtubule assay used to examine the collective activity of Kif4A and CLASP1 on single microtubules. HiLyte 647 labeled seeds (blue) were immobilized on glass coverslips via a neutravidin-biotin linkage. The seeds were incubated with X-Rhodamine labeled tubulin, CLASP1-GFP and Kif4A and imaged for 20 minutes. Polarity of microtubules are indicated as + and -.

**b**. Scatter plot with bar graph showing mean of fraction of time microtubules are stalled. Error bars indicate standard deviation. Assay conditions: tubulin control (0.0 ± 0.0, n = 59), 200 nM CLASP1 (0.0 ± 0.0, n = 67), 200 nM CLASP1 + 125 nM Kif4A (0.3 ± 0.3, n = 67), 200 nM CLASP1 + 250 nM Kif4A (0.4 ± 0.3, n = 73), 200 nM CLASP1 + 1000 nM Kif4A (0.6 ± 0.4, n = 73) and 1000 nM Kif4A (0.9 ± 0.1, n = 39).

**c.** Box and whisker plot of average length of microtubule before catastrophe. Plus-sign indicates mean. Horizontal lines within box indicate the 25th, median (center line) and 75th percentile. Error bars indicate minimum and maximum range. Mean and standard deviation for assay conditions: Tubulin control (2.9 ± 2.4 µm, n = 241), 200 nM CLASP1 (6.1 ± 3.2 µm, n = 150), 200 nM CLASP1 + 125 nM Kif4A (4.0 ± 1.9 µm, n = 82), 200 nM CLASP1 + 250 nM Kif4A (3.2 ± 1.8 µm, n = 107), 200 nM CLASP1 + 1000 nM Kif4A (1.7 ± 1.0 µm, n = 152) and 1000 nM Kif4A (0.9 ± 0.6 µm, n = 24). P < <0.0001 for 1000 nM Kif4A compared to tubulin control.

**d.** Scatter plot of Kif4A-GFP intensity per pixel on taxol-stabilized microtubules in the presence of 150 µM ATP (n = 70 microtubules from 2 independent experiments). Concentrations refer to monomeric Kif4A. Mean and standard deviation of intensity for assay conditions with Kif4A-GFP: 20 nM (504.2 ± 395.8, n=54), 100 nM (2556 ± 1182, n=69), 400 nM (7823 ± 1961, n=68), 1000 nM (8276 ± 3386, n=69).

**e.** Box and whisker plot of average number of catastrophes in 20 minutes. Plus-sign indicates mean. Horizontal lines within box indicate the 25th, median (center line) and 75th percentile. Error bars indicate minimum and maximum range. Mean and standard deviation for assay conditions: tubulin control (5.1 ± 2.2, n = 65), 200 nM CLASP1 (1.9 ± 1.6, n = 76), 200 nM CLASP1 + 125 nM Kif4A (1.5 ± 1.5, n = 69), 200 nM CLASP1 + 250 nM Kif4A (1.8 ± 1.7, n = 73), 200 nM CLASP1 + 1000 nM Kif4A (2.3 ± 2.5, n = 73) and 1000 nM Kif4A (1.6 ± 3.2, n = 38). P < 0.0001 for (i) tubulin control to 200 nM CLASP1. P is not significant (> 0.05) for 200 nM CLASP1 to (i) 200 nM CLASP1 + 125 nM Kif4A, (ii) 200 nM CLASP1 + 250 nM Kif4A, (iii) 200 nM CLASP1 + 1000 nM Kif4A and to (iv) 1000 nM Kif4A.

**f.** Box and whisker plot of average number of rescues in 20 minutes. Plus-sign indicates mean. Horizontal lines within box indicate the 25th, median (center line) and 75th percentile. Error bars indicate minimum and maximum range. Mean and standard deviation for assay conditions: tubulin control (0.1 ± 0.4, n = 60), 200 nM CLASP1 (2.0 ± 1.5, n = 67), 200 nM CLASP1 + 125 nM Kif4A (1.4 ± 1.5, n = 67), 200 nM CLASP1 + 250 nM Kif4A (1.4 ± 1.5, n = 73), 200 nM CLASP1 + 1000 nM Kif4A (0.9 ± 1.3, n = 73) and 1000 nM Kif4A (0.0 ± 0.0, n = 37). P < 0.0001 for tubulin control to 200 nM CLASP1 and P = 0.0025 for 200 nM CLASP1 + 1000 nM Kif4A to 1000 nM Kif4A.

**g.** Scatter plot of CLASP1-GFP fluorescence intensity per pixel on single microtubules in the presence of CLASP1-GFP and Kif4A at indicated concentrations. Red bars indicate mean (center line) and standard error of mean. Assay conditions: 200 nM CLASP1 (25.9 ± 4.6, n = 59), 200 nM CLASP1 + 125 nM Kif4A (23.0 ± 6.5, n = 59), 200 nM CLASP1 + 250 nM Kif4A (18.4 ± 6.3, n = 60) and 200 nM CLASP1 + 1000 nM Kif4A (12.8 ± 6.3, n = 59). P values < 0.0001 for 200 nM CLASP1 when compared to (i) 200 nM CLASP1 + 250 nM Kif4A and (ii) 200 nM CLASP1 and 1000 nM Kif4A. n is number of microtubules analyzed from 3 independent experiments.

### Extended Data Fig. 3

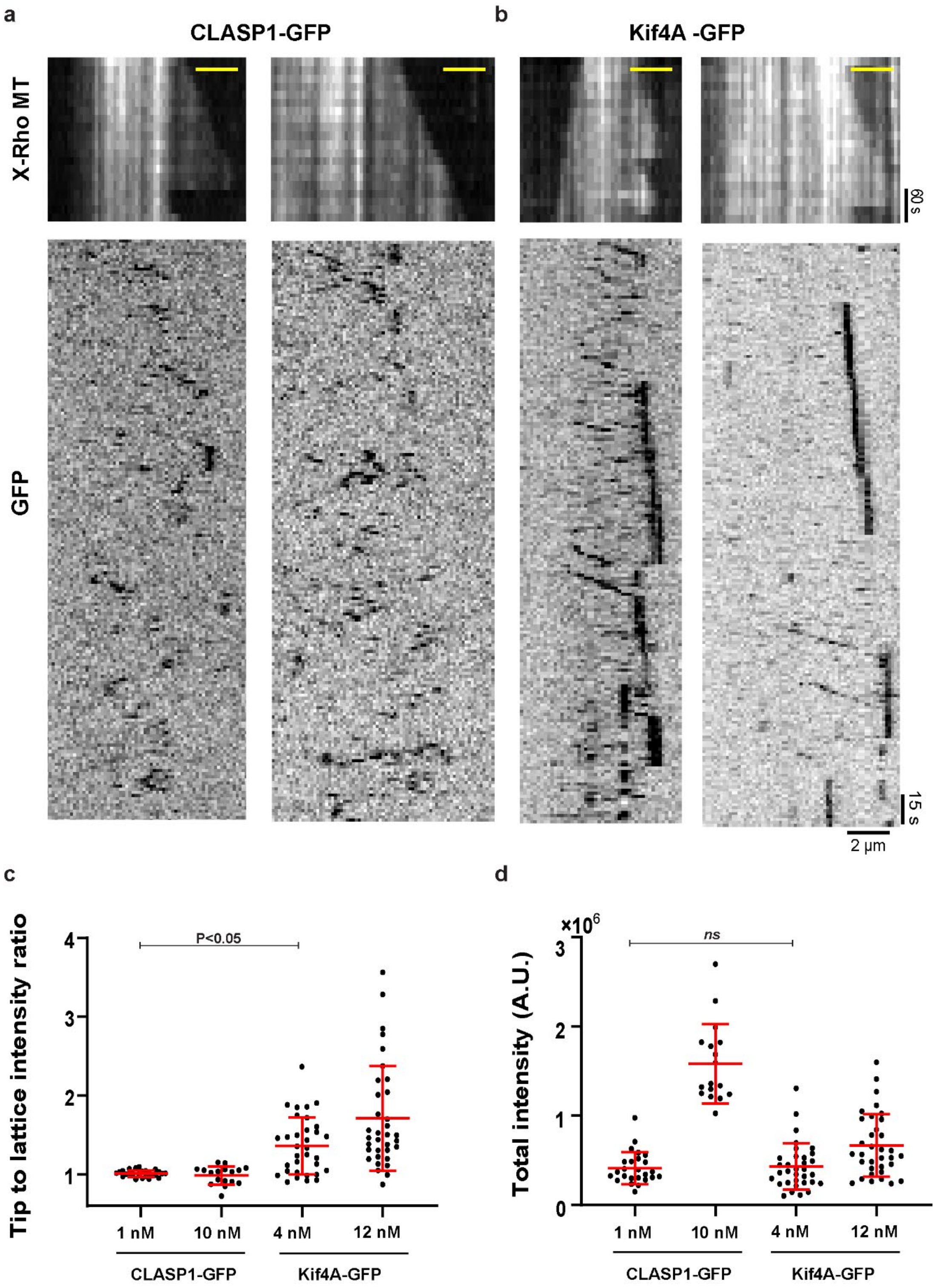


**Extended Data Fig. 3.** *Related to Fig. 1*

n = number of kymographs analyzed from 3 independent experiments.

P values were calculated from an ordinary one-way ANOVA test with Dunnett correction for multiple comparisons.

All concentrations refer to monomeric GFP-tagged proteins.

**a.** Representative kymographs generated from single molecule TIRF experiments of 1 nM CLASP1-GFP on dynamic microtubules, showing X-rhodamine microtubule (Top, X-Rho MT) and GFP (bottom) channels, from 25 kymographs analyzed. Scale bars for X-Rho MT channel: x=2 μm, y=60 s; for GFP channel: x= 2 μm and y=15 s.

**b.** Representative kymographs generated from single molecule TIRF experiments of 12 nM Kif4A-GFP. on dynamic microtubules, showing X-rhodamine microtubule (Top, X-Rho MT) and GFP (bottom) channels from 34 kymographs analyzed. Scale bars for X-Rho MT channel: x=2 μm, y=60 s; for GFP channel: x= 2 μm and y=15 s.

**c.** Scatter plot of tip to lattice GFP intensity ratio. Horizontal line indicates the mean. Error bars indicate standard deviation. Tip to lattice intensity ratios for 1 nM CLASP1-GFP (1.01 ± 0.04; n =25); 10 nM CLASP1-GFP (0.98 ± 0.11; n =17); 4 nM Kif4A-GFP (1.36 ± 0.36; n =33); 12 nM Kif4A-GFP (1.71 ± 0.66; n =34). P = 0.0066 for 1 nM CLASP1 compared to 4 nM Kif4A.

**d.** Scatter plot of the total GFP intensity. Horizontal line indicates the mean. Error bars indicate standard deviation. Total intensity values for 1 nM CLASP1-GFP ($4.14 \times 10^5 \pm 1.80 \times 10^5$; n = 25); 10 nM CLASP1-GFP ($1.58 \times 10^5 \pm 4.43 \times 10^4$; n = 17); 4 nM Kif4A-GFP ($4.31 \times 10^5 \pm 2.60 \times 10^5$; n=33); 12 nM Kif4A-GFP ($6.66 \times 10^5 \pm 3.50 \times 10^5$; n =34). P is not significant for for 1 nM CLASP1 compared to 4 nM Kif4A.

### Extended Data Fig. 4

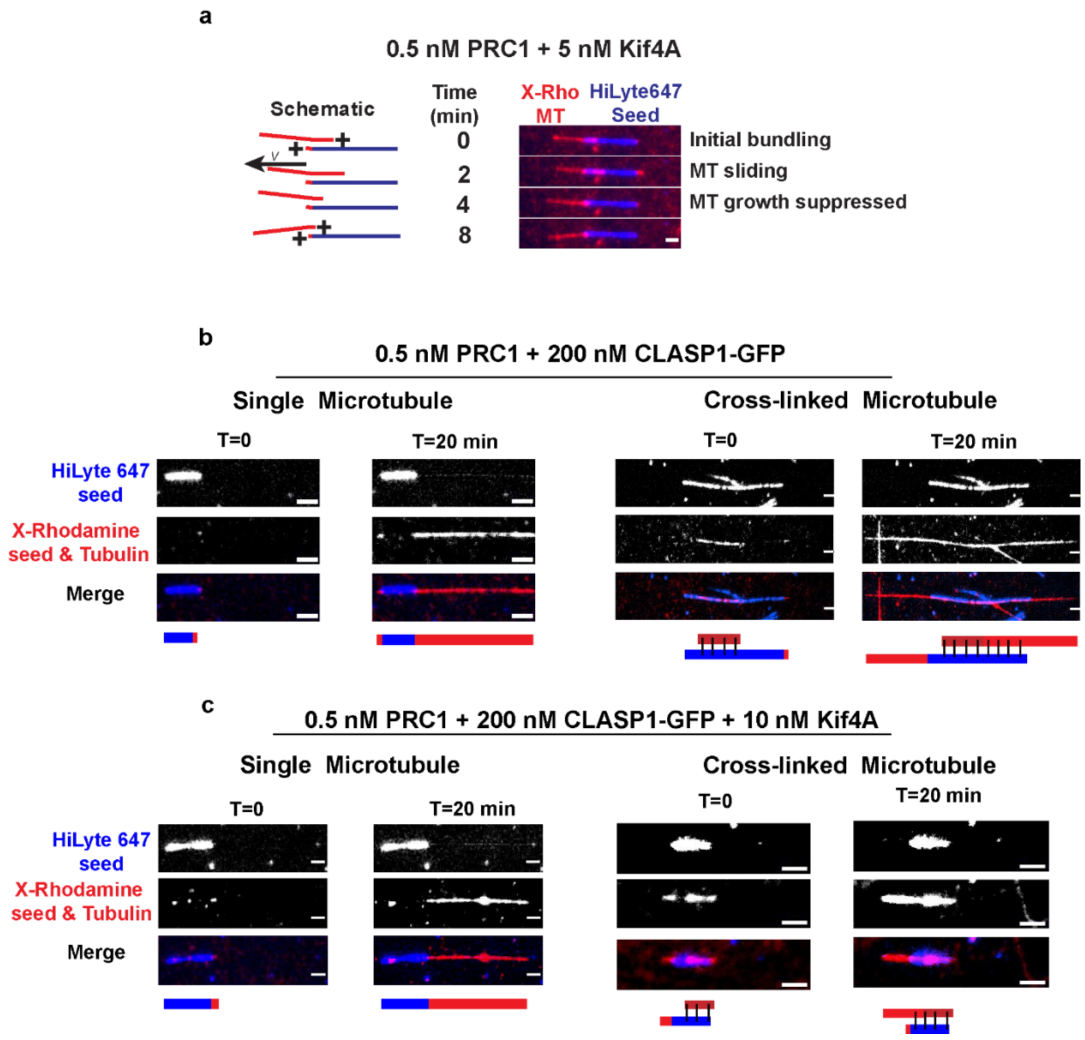


**Extended Data Fig. 4.** *Related to Fig. 2*

**a.** Schematics and montages of representative microtubule bundles (red) grown from microtubule seeds (blue) in the presence of 5 nM Kif4A and 0.5 nM PRC1. A total of 20 events were examined. Schematics indicate the plus end (+) of the microtubules within the bundle. Velocity arrow indicates direction of microtubule sliding. X-Rh MT: X-Rhodamine microtubules. Scale bar represents 2 μm.

**b & c.** Representative image of single microtubule (left) and cross-linked microtubule (right), showing HiLyte 647 (top), X-Rhodamine (middle) and merged (bottom) channels at the start (T=0) and end (T=20 min) of dynamic bundle assay. Scale bars represent 2 μm. The schematics below the montages indicate the positions of the seed (blue), microtubules (red) at the start and end of the experiment. Assay conditions are **(b)** 0.5 nM PRC1 + 200 nM CLASP1-GFP (Events examined: 47 single and 29 overlaps), and **(c)** 0.5 nM PRC1 + 200 nM CLASP1-GFP + 10 nM Kif4A (Events examined: 42 single and 49 overlaps).

### Extended Data Fig. 5

**a**

| | | | | | | | |
|---|---|---|---|---|---|---|---|
| CLASP1 (nM) | 200 | 200 | 200 | 200 | 200 | 20 | 0 |
| Kif4A (nM) | 125 | 50 | 10 | 5 | 0 | 5 | 5 |
| PRC1 (nM) | 0.5 | 0.5 | 0.5 | 0.5 | 0.5 | 0.5 | 0.5 |
| Dynamic Instability (%) | 0 | 0 | 0 | 56 | 86 | 90 | 70 |
| No Growth (%) | 100 | 70 | 78 | 0 | 0 | 8 | 15 |
| Only Growth (%) | 0 | 30 | 22 | 44 | 14 | 2 | 15 |
| Total Number of Events | 30 | 33 | 36 | 50 | 63 | 61 | 53 |
| Complete Catastophes (%) | 0 | 0 | 0 | 7 | 2 | 64 | 100 |
| Rescue (%) | 0 | 0 | 0 | 93 | 98 | 36 | 0 |

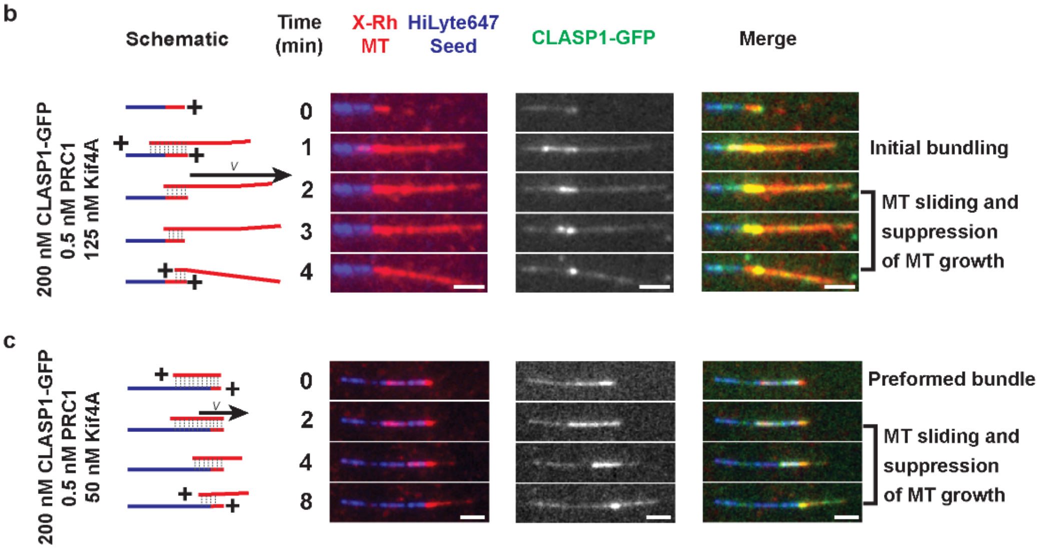


**Extended Data Fig. 5.** *Related to Fig. 3*

**a.** Summary of dynamics of PRC1 cross-linked microtubules, showing percentage of microtubules exhibiting dynamic instability, no growth or only growth in black. For all kymographs, the total number of events, percentage of events that are complete catastrophes and rescues in the presence of CLASP1-GFP, PRC1 and Kif4A at varying concentrations are shown in blue.

**b & c.** Schematics and representative montages of microtubule bundles (red) grown from microtubule seeds (blue). Schematics indicate the plus end of the microtubules within the bundle. Velocity arrow indicates direction of microtubule sliding. Dotted gray lines indicate regions of overlap. X-Rh MT: X-Rhodamine microtubules. Scale bar represents 2 µm. Assay conditions and number of events (n) examined across 3 independent experiments are (**b)** 200 nM CLASP1-GFP + 0.5 nM PRC1 + 125 nM Kif4A (n=30) and **(c)** 200 nM CLASP1-GFP + 0.5 nM PRC1 + 50 nM Kif4A (n=39).

**Extended Data Fig. 6**

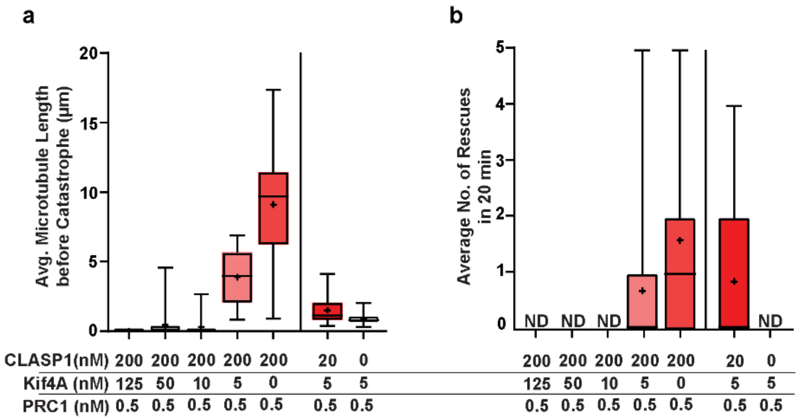


**Extended Data Fig. 6.** *Related to Fig. 4*

n = number of kymographs of cross-linked microtubules analyzed

**a.** Box and whisker plot of average microtubule length before catastrophe of cross-linked microtubules. Plus-sign indicates mean. Horizontal lines within box indicate the 25th, median (center line) and 75th percentile. Error bars indicate minimum and maximum range. Mean and standard deviation for assay conditions: 200 nM CLASP1 + 0.5 nM PRC1 + 125 nM Kif4A (0.0 ± 0.0 nm, n = 30), 200 nM CLASP1 + 0.5 nM PRC1 + 50 nM Kif4A (0.4 ± 0.9 nm, n = 39), 200 nM CLASP1 + 0.5 nM PRC1 + 10 nM Kif4A (0.3 ± 0.7 nm, n = 36), 200 nM CLASP1 + 0.5 nM PRC1 + 5 nM Kif4A (3.9 ± 1.9 nm, n = 28), 200 nM + 0.5 nM PRC1 (9.1 ± 3.6 nm, n = 53), 20 nM CLASP1 + 0.5 nM PRC1 + 5 nM Kif4A (1.5 ± 0.9 nm, n = 55) and 0.5 nM PRC1 + 5 nM Kif4A (0.9 ± 0.4 nm, n = 37).

**c.** Box and whisker plot of average microtubule length before catastrophe of cross-linked microtubules. Plus-sign indicates mean. Horizontal lines within box indicate the 25th, median (center line) and 75th percentile. Error bars indicate minimum and maximum range. Mean and standard deviation for assay conditions 200 nM CLASP1 + 0.5 nM PRC1 + 125 nM Kif4A (*Not Determined*, n = 30), 200 nM CLASP1 + 0.5 nM PRC1 + 50 nM Kif4A (*Not Determined*, n = 39), 200 nM CLASP1 + 0.5 nM PRC1 + 10 nM Kif4A (*Not Determined*, n = 36), 200 nM CLASP1 + 0.5 nM PRC1 + 5 nM Kif4A (0.7 ± 1.2, n = 37), 200 nM + 0.5 nM PRC1 (1.6 ± 1.5, n = 33), 20 nM CLASP1 + 0.5 nM PRC1 + 5 nM Kif4A (0.9 ± 1.3, n = 23) and 0.5 nM PRC1 + 5 nM Kif4A (0.0 ± 0.0, n = 20).

**Extended Data Fig. 7.**

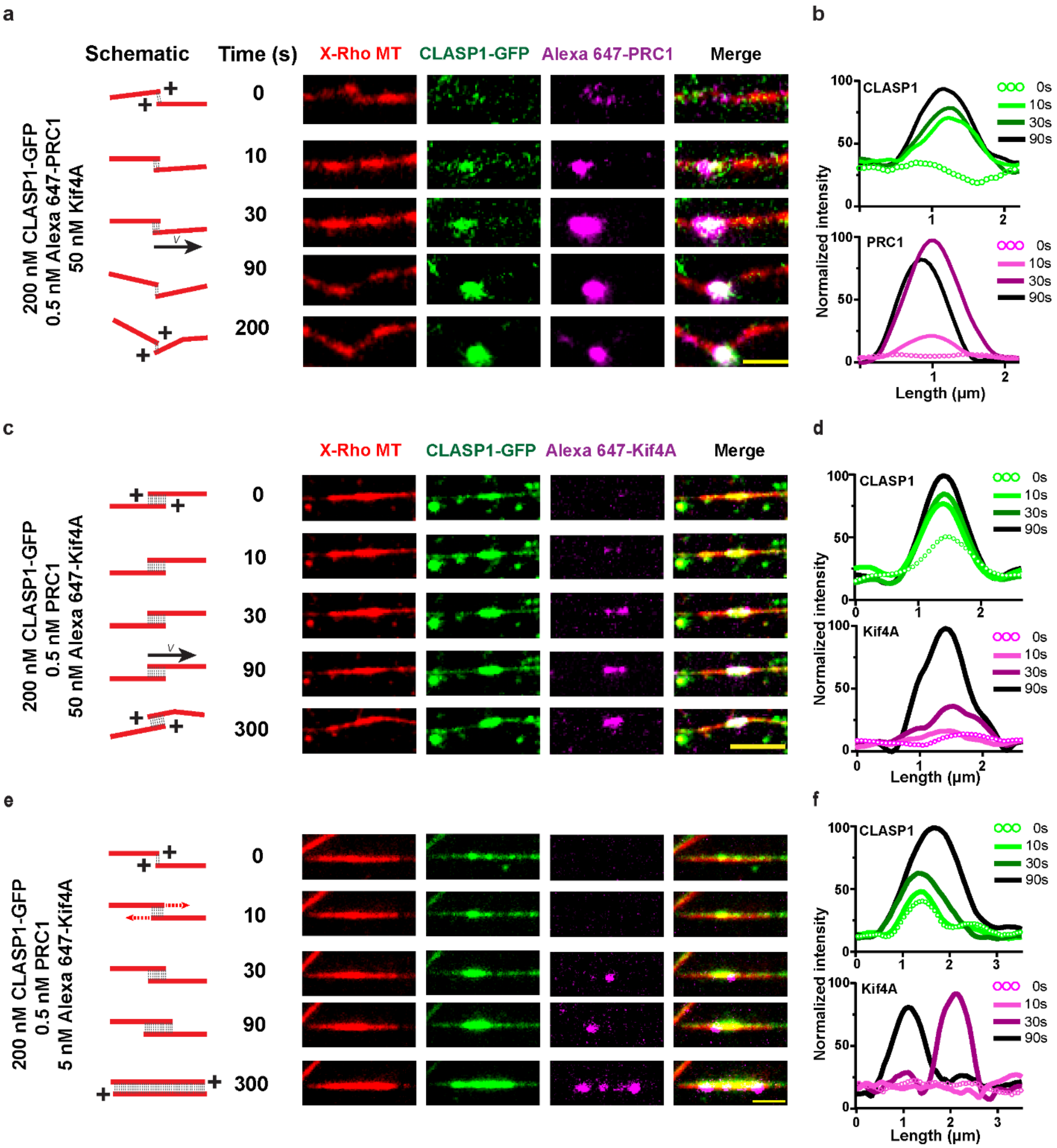


**Extended Data Fig. 7.** *Related to Fig. 5*

**a, c, e.** Schematics and montages of X-rhodamine labeled microtubules (red) which grow from X-rhodamine labeled seeds (red) and form new bundles. Time T= 0 refers to the first image where the two growing ends encounter each other, as judged by the intensity in the tubulin channel. Schematics indicate the plus end of the microtubules within the bundle. Velocity arrow indicates direction of microtubule sliding. Dotted red arrows indicate microtubule growth. Dotted gray lines indicate regions of overlap. X-Rho MT: X-Rhodamine microtubules. Scale bar represents 2 μm.

**b, c, f.** Line scans of intensities of GFP (top) and Alexa-647 (bottom) channels along a line joining the two microtubules and containing the overlap, at times T = 0, 10s 30s and 90s following bundle formation. Intensities were normalized between 0 (0 intensity) and 100 for the maximum intensity recorded for each channel, across all time points.

Assay conditions are

**a & b:** 200 nM CLASP1-GFP + 0.5 nM Alexa 647-labeled PRC1 + 50 nM Kif4A (8 events analyzed from 2 independent experiments)

**c & d:** 200 nM CLASP1-GFP + 0.5 nM PRC1 + 50 nM Alexa 647-labeled Kif4A (31 events analyzed from 3 independent experiments)

**e & f:** 200 nM CLASP1-GFP + 0.5 nM PRC1 + 5 nM Alexa 647-labeled Kif4A (22 events analyzed from 3 independent experiments)

### Extended Data Fig. 8

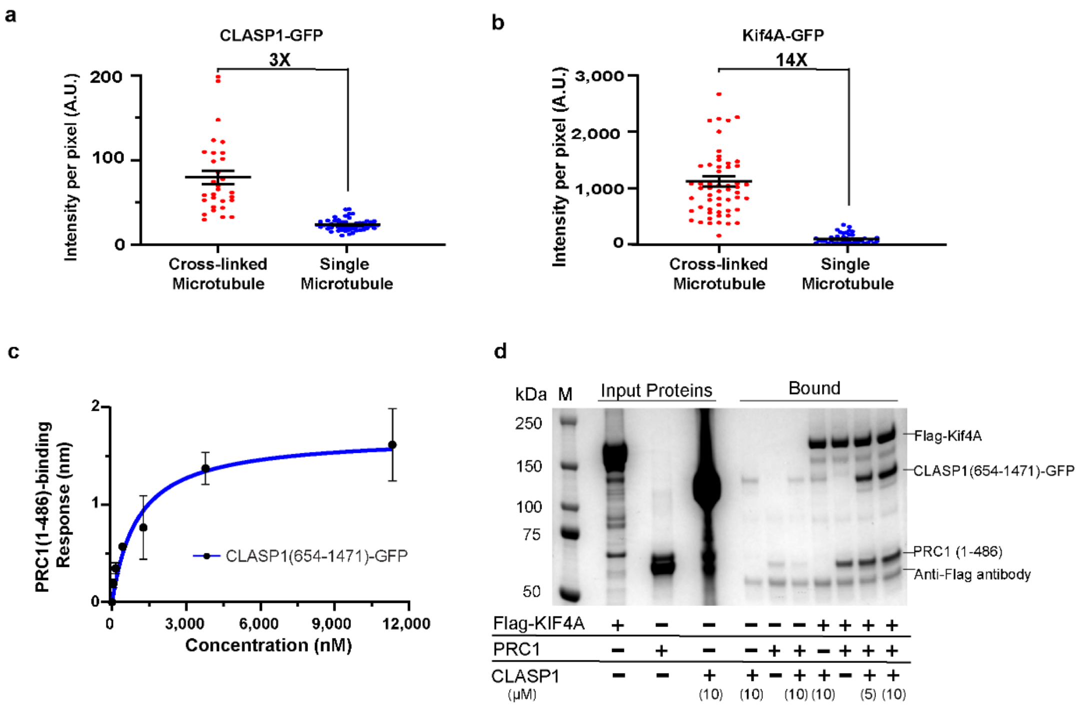


**Extended Data Fig. 8.** *Related to Fig. 5*

**a.** Scatter plot of CLASP1-GFP fluorescence intensities on single (•) and cross-linked (•) microtubules. Mean intensity and standard error of mean for assay conditions: 0.5 nM PRC1 + 200 nM CLASP1-GFP, single microtubules (24.2 ± 0.9, n=54), cross-linked microtubules (80.2 ± 8.1, n=30). n = number of microtubules analyzed from 3 independent experiments in each condition.

**b.** Scatter plot of Kif4A-GFP fluorescence intensities on single (•) and cross-linked (•) microtubules. Mean intensity and standard error of mean for assay conditions: 0.5 nM PRC1 + 10 nM Kif4A-GFP, single microtubules (80.1 ± 13, n=46), cross-linked microtubules (1106 ± 90.1, n=57). n = number of microtubules analyzed from 3 independent experiments in each condition.

**c.** BLI assay to quantify the binding affinity of CLASP1(654-1471)-GFP to PRC1(1-486). Error bars represent standard error of mean. Data from 3 independent experiments were fit to a Hill equation. $K_D$ : 1.04 ± 0.44 μM ($R^2$ of fit = 0.90).

**d.** Representative SDS-PAGE gel showing input proteins (lanes 2-4) and proteins appearing in the bound fraction (lanes 5-11) of pull-down assay with immobilized Flag-Kif4A, from one of 3 independent experiments. Reaction conditions corresponding to each lane are described in the table below the gel. Concentration of CLASP1(654-1471)-GFP used in each reaction is indicated in parentheses below the table. M: molecular weight marker, with corresponding masses in kDa on the side.

### Extended Data Fig. 9

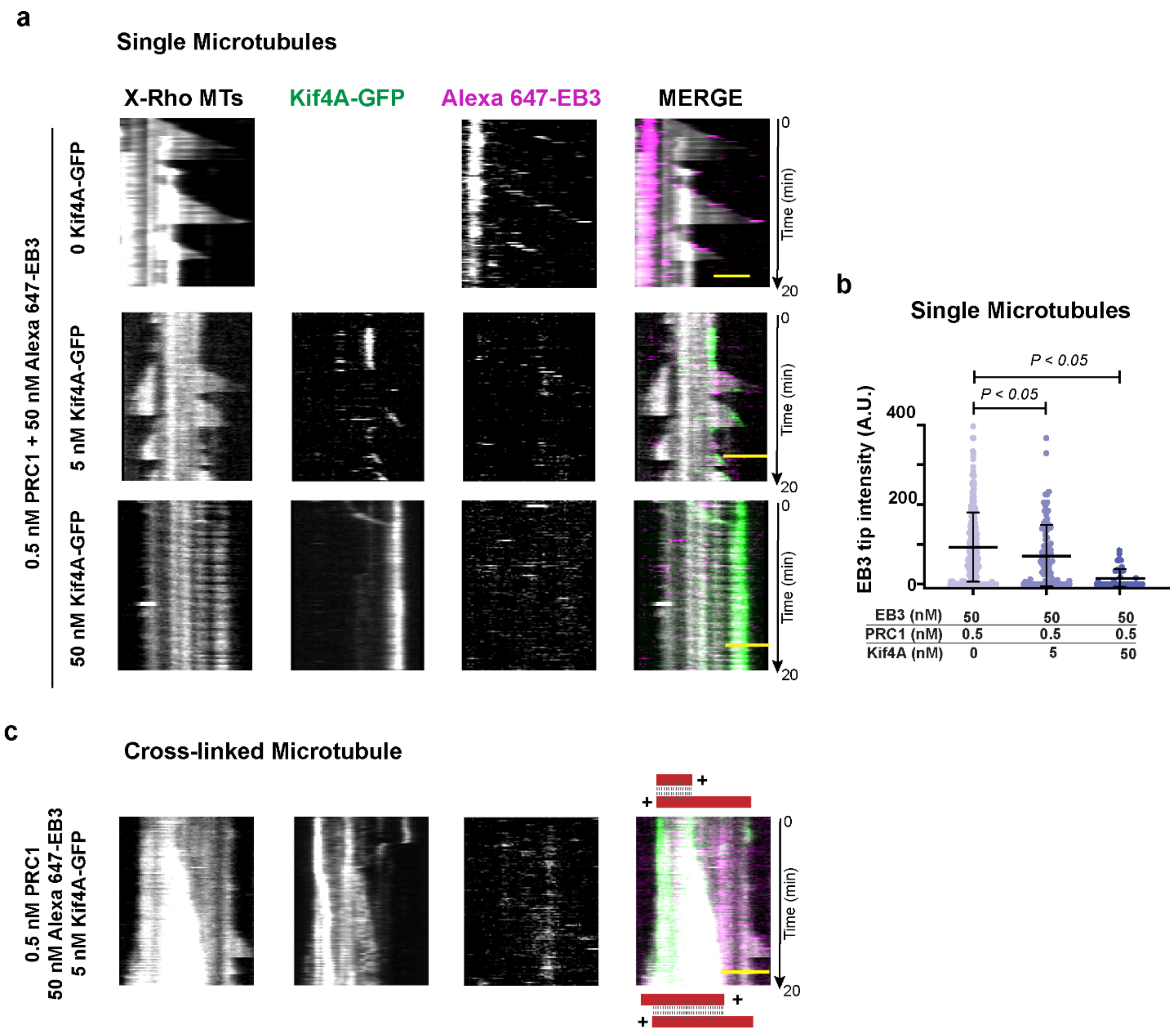


**Extended Data Figure 9.** *Related to Fig. 5*

**a.** Representative kymographs of dynamic single X-rhodamine-labeled microtubules (X-Rho MTs) with 50 nM Alexa 647-labeled EB3 + 0.5 nM PRC1 (n = 70), 50 nM Alexa 647-labeled EB3 + 0.5 nM PRC1 + 5 nM Kif4A-GFP (n = 93), 50 nM Alexa 647-labeled EB3 + 0.5 nM PRC1 + 50 nM Kif4A-GFP (n = 60). Scale bar represents 2 μm. n = number of Kymographs analyzed from 3 independent experiments

**b.** Scatter plot of intensities of EB3 at microtubule tips with increasing concentrations of Kif4A-GFP. Mean and standard deviation are indicated by black bars. Assay condition (n=number of growth events analyzed): 50 nM Alexa 647-labeled EB3 + 0.5 nM PRC1 (186.40 ± 173.80, n=281), 50 nM Alexa 647-labeled EB3 + 0.5 nM PRC1 + 5 nM Kif4A-GFP (143.80 ± 155.20, n=98), 50 nM Alexa 647-labeled EB3 +

0.5 nM PRC1 + 50 nM Kif4A-GFP (28.11 ± 47.85, n=43). For 50 nM Alexa 647-labeled EB3 + 0.5 nM PRC1compared to (i) 50 nM Alexa 647-labeled EB3 + 0.5 nM PRC1 + 5 nM Kif4A-GFP, P= 0.048, (ii) 50 nM Alexa 647-labeled EB3 + 0.5 nM PRC1 + 50 nM Kif4A-GFP, P < <0.0001, in an ordinary one-way ANOVA test with Dunnett correction for multiple comparisons.

**c.** Representative kymographs of a cross-linked X-rhodamine-labeled microtubule bundle in the presence of 50 nM Alexa 647-labeled EB3, 0.5 nM PRC1 and 5 nM Kif4A-GFP. Schematics above and below the merged kymograph indicate positions of the microtubules (red) and PRC1 (gray dashed line) at the start and end of the experiment, respectively. + indicates microtubule polarity. Scale bar represents 2 μm. A total of 45 kymographs from 3 independent experiments were examined.

**Extended Data Fig. 10**

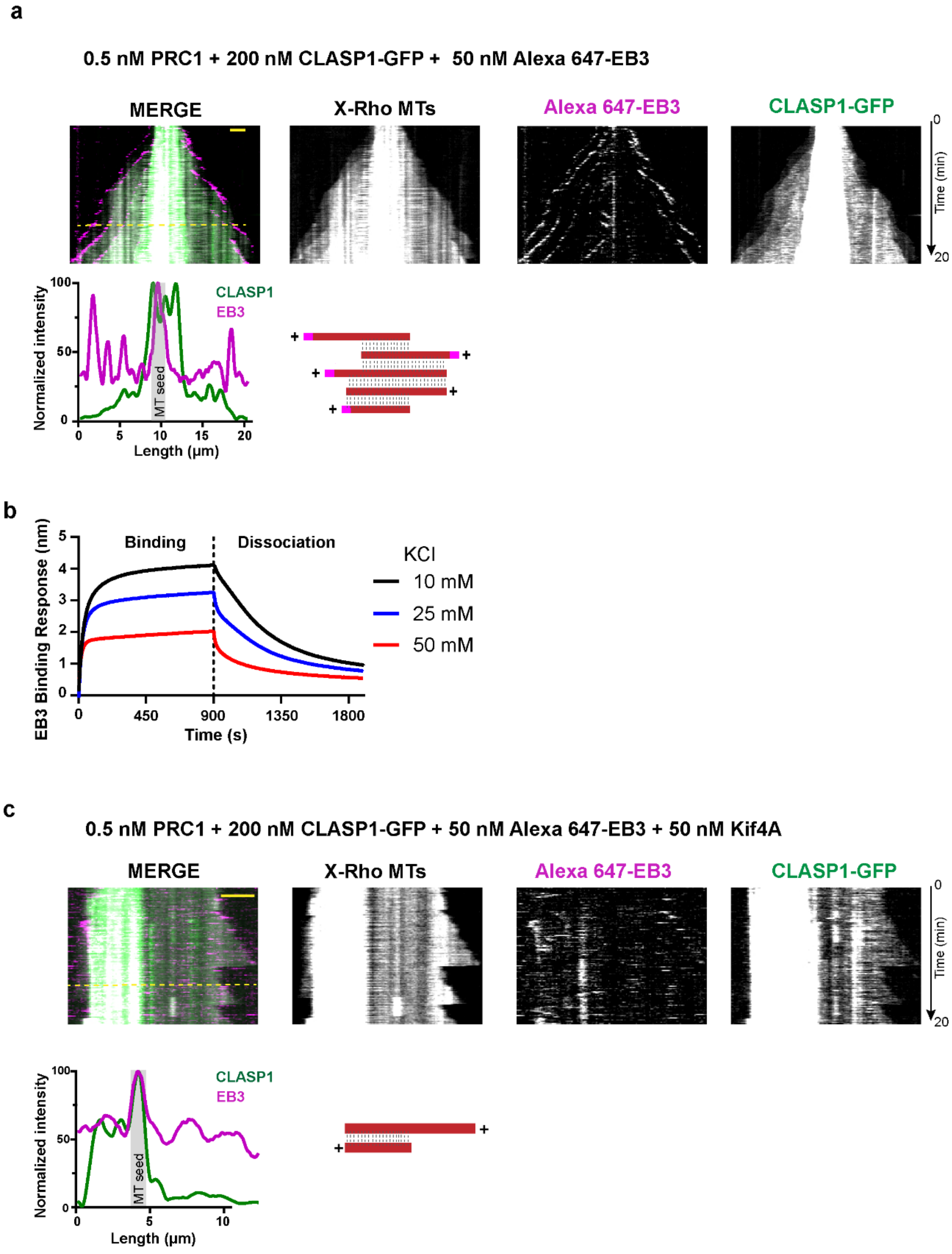


**Extended Data Figure 10.** *Related to Fig. 5*

**a.** Representative kymographs of cross-linked X-rhodamine-labeled microtubule bundle (X-Rho MTs) in the presence of 0.5 nM PRC1, 200 nM CLASP1-GFP and 50 nM Alexa 647-labeled EB3. Intensity profile below merged kymograph shows the normalized intensities of the GFP (green) and Alexa 647 (magenta) channels at the time-point represented by dashed yellow line on kymograph. Gray box indicates the position of microtubule seeds. Intensities were normalized between 0 (0 intensity) and 100 for the maximum intensity recorded for each channel. Schematics on bottom right represent the positions of microtubules (red), EB3 (magenta) and PRC1 (gray dashed lines) at the same time-point. Scale bar represents 2 μm.  A total of 73 kymographs from 3 independent experiments were examined.

**b.** Representative BLI sensorgram of the binding response of 1μM CLASP1(654-1471)-GFP in solution to immobilized EB3, in the presence of 10 mM (black), 25 mM (blue) and 50 mM (red) KCl. Vertical dotted black line demarcates the binding and dissociation phases. The sensorgrams have been corrected for drift of EB3 from the sensor and non-specific binding of CLASP1(654-1471)-GFP to the sensor. Binding curves from 3 independent experiments were analyzed.

**c.** Representative kymographs of cross-linked X-rhodamine-labeled microtubules (X-Rho MTs) in the presence of 0.5 nM PRC1, 200 nM CLASP1-GFP, 50 nM Kif4A and 50 nM Alexa 647-labeled EB3. Intensity profile below merged kymograph shows the normalized intensities of the GFP (green) and Alexa-647 (magenta) channels at the time-point represented by dashed yellow line on kymograph. Gray box indicates the position of microtubule seeds. Intensities were normalized between 0 (0 intensity) and 100 for the maximum intensity recorded for each channel. Schematics on bottom right represent the positions of microtubules (red), EB3 (magenta) and PRC1 (gray dashed lines) at the same time-point. Scale bar represents 2 μm. A total of 67 kymographs from 3 independent experiments were examined.
